# Central infusion of prostaglandin E2 reveals a unified representation of sickness in the mouse insular cortex

**DOI:** 10.1101/2025.04.28.651028

**Authors:** Gretel B. Kamm, Juan C. Boffi, Muad Y. Abd El Hay, Domitille Rajot, Ana Cukić, Martha N. Havenith, Marieke Scholvinck, Nicolas Renier, Hiroki Asari, Robert Prevedel

**Affiliations:** Cell Biology and Biophysics Unit, European Molecular Biology Laboratory, Heidelberg, Germany; Zero-Noise Lab, Ernst Strüngmann Institute For Neuroscience, Frankfurt Am Main, Germany; Sorbonne Université, Paris Brain Institute, INSERM, CNRS, AP-HP, Paris, France; Epigenetics and Neurobiology Unit, European Molecular Biology Laboratory, Rome, Italy; Interdisciplinary Center for Neurosciences, Heidelberg University, Heidelberg, Germany

**Keywords:** Sickness Syndrome, Prostaglandin E2, Insular Cortex, Primary Somatosensory Cortex

## Abstract

During infections, vertebrates develop stereotypic symptoms such as elevated body temperature, reduced appetite, and lethargy. These changes, collectively known as sickness syndrome, are orchestrated by the brain in response to immune mediators released during systemic inflammation. While the roles of subcortical regions, including the hypothalamus and brainstem nuclei, in regulating sickness symptoms are well established, the contribution of the neocortex to the encoding and modulation of the sick state remains less well understood. We examined the neuronal correlates of sickness in the neocortex of awake mice following a single intracerebroventricular (i.c.v.) injection of prostaglandin E2 (PGE2), a well-characterized mediator of sickness. Behavioral analysis revealed that PGE2 elicited a rapid and robust sickness response, characterized by fever, slower locomotion, quiescence, anorexia, and eye squinting. Whole-brain Fos mapping showed that PGE2 generates a distinct neural activation pattern encompassing much of the interoceptive network. Electrophysiological recordings using Neuropixel probes in awake mice together with dimensionality reduction and decoding analysis revealed that neuronal population dynamics in the insular cortex (IC) and the primary somatosensory cortex (SSp), two regions involved in body state representation, encode sickness-related information, such as body temperature, walking velocity, grooming, and eye squinting. However, unlike SSp, ongoing neuronal activity in IC exhibited a better decoding performance for an integrated measure of sickness rather than individual symptoms. Together, these results suggest that PGE2 induces a coordinated physiological and behavioral response akin to a sick state, which is preferentially encoded in the IC.

## Introduction

Infections often induce profound changes in both behavior and physiology in vertebrates, including humans. These changes can manifest as behavioral symptoms, such as decreased locomotion and reduced social and feeding behaviors. In addition, infections trigger autonomic and endocrine adaptations, including increased body temperature and corticosterone/cortisol release^1–6^. The brain coordinates these changes, collectively known as sickness syndrome, to prepare the organism for fighting infections, thereby improving survival chances for the individual and their group^7–9^.

The sickness response begins when the immune system detects pathogen-associated molecules. In response, the immune system releases a complex mixture of inflammatory mediators. These mediators then inform the brain about the impending infection by targeting nerve endings in peripheral tissues or by reaching the brain directly through the bloodstream^9^.

Studies in humans^10,11^ and mice^1–3,12–14^ have shown that multiple brain regions respond to peripheral inflammation. Interestingly, many of these inflammation-responsive areas are also involved in processing information about the internal conditions of the body, a process known as interoception^15^. This suggests that inflammation may trigger a distinct bodily state representation in the brain^10,16,17^, which could be identified as a “sick state”.

While the neuronal correlates of physiological states such as hunger or thirst have been extensively studied^18–21^, much less is known about how the brain encodes the sick state during infections, a process recently termed immunoception^16^, at a single neuron resolution. One limitation for further progress is that the most common method to trigger sickness, a single injection of the gram-negative bacterial cell wall component lipopolysaccharide (LPS)^22^, produces a slow sick state development that can last hours^1–3^. The lengthy and polyphasic nature of LPS-triggered sickness^23^ makes it challenging to implement during neurophysiological recordings under non-anesthetized conditions.

An alternative approach to study sickness representation in awake animals relies on intracerebroventricular (i.c.v.) injections of fast-acting immune mediators. Among the best-studied direct mediators of sickness is prostaglandin E2 (PGE2)^24^. PGE2 is a short-lived lipid derived from arachidonic acid that has roles in diverse processes, including reproduction, metabolism regulation, and immunity^25,26^. During immune challenges, PGE2 production is highly induced in endothelial cells within the brain^27,28^. Neurons can detect PGE2 directly through four specific G protein-coupled receptors^29^ expressed across multiple brain areas^30–33^. Acting directly in the brain, PGE2 can trigger many sick-associated symptoms such as fever^33^, increased pain sensitivity^34,35^, warmth-seeking behaviors^36^, and loss of appetite^37,38^.

Combining behavioral analysis, whole-brain activity mapping, and *in vivo* electrophysiology, we studied how neuronal activity changes during sickness. We observed that PGE2 induces a rapid sickness response in mice marked by high fever, slower locomotion, quiescence, anorexia, and eye squinting. Furthermore, PGE2 activates brain areas involved in thermoregulation, energy homeostasis, and stress. Neuropixel recordings in awake mice combined with state-of-the-art dimensionality reduction and decoding approaches revealed that the insular cortex (IC) encodes an integrated representation of sickness significantly better than the primary somatosensory cortex (SSp). These findings establish central PGE2 injections as a valuable tool for studying the neural representation of sickness in awake mice and highlight a potential role of the IC in representing immune system states.

## Results

### A central injection of PGE2 generates a robust sickness response in mice

Systemic administration of the bacterial cell-wall component LPS generates a comprehensive sickness response in mice^1,2,39^ by activating the peripheral immune system and triggering the production of intermediates such as prostaglandins^9,28^. Prostaglandins are widely regarded as an essential component of the sickness response, and blocking their production is sufficient to ameliorate many of the symptoms associated with sickness^8^. This constitutes the primary mechanism of action for drugs, such as acetylsalicylic acid (aspirin) and ibuprofen^26^. Given the centrality of prostaglandins in sickness^8^ and their quick turnover^26^, we explored the use of brain injections of PGE2 as a method to trigger a fast and short-lived sickness response in mice.

Combining stress-free body temperature measurements, automated tools for estimating mouse postures^40^ and supervised machine learning pipelines for classifying behaviors^41^, we obtained a detailed profile of the sickness responses elicited by peripherally administered LPS and by centrally infused PGE2 in mice (**Figure 1A and B**). This approach allowed us to accurately extract behavioral events in home-caged mice, even those that were partially occluded, such as interacting with food pellets under the food tray (**Figure S1A and B**).

**Figure 1:**
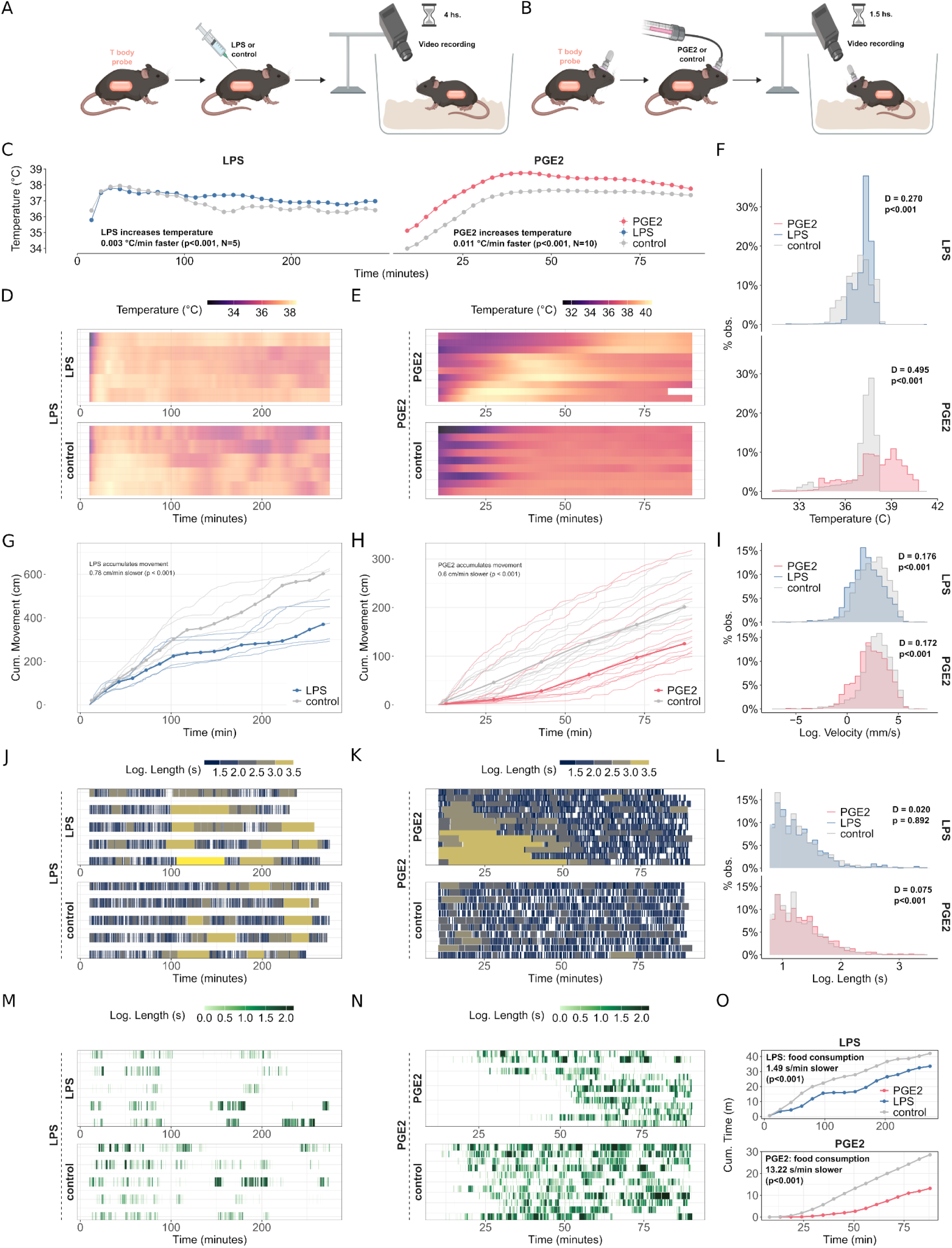
A central injection of PGE2 generates a robust sickness response in mice. **(A-B)** Schematic illustrations of the experimental approach to study sickness in freely-moving mice upon (A) an i.p. injection of LPS or (B) i.c.v. injection of PGE2. **(C)** Mean body temperature profiles over time in PGE2 and LPS experiments compared to their respective controls (linear mixed models with time as a factor). **(D-E)** Temperature heatmaps for individual mice in LPS and PGE2 experiments, arranged by temperature response magnitude. **(F)** Temperature distribution histograms for both experimental groups (Kolmogorov-Smirnov tests). **(G-H)** Median cumulative movement over time in LPS- and PGE2-treated mice compared to controls (linear mixed models with time as a factor). **(I)** Velocity distribution histograms (Kolmogorov-Smirnov tests). **(J-K)** Stasis episode visualization for LPS and PGE2 experiments, where each horizontal segment represents an immobility episode, with color intensity indicating the logarithm of episode duration. Animals were ordered by stasis episode length and frequency. **(L)** Distribution of stasis episode durations (log scale) for PGE2 and LPS experiments (Kolmogorov-Smirnov tests). **(M-N)** Food interaction episodes for LPS and PGE2 experiments, with each horizontal segment representing a feeding episode and color indicating the logarithm of episode duration. Animals were ordered in the same order as figures J and K. **(O)** Mean cumulative time spent on food interaction over time for treatment and control groups (linear mixed models with time as a factor). See **Table 1** for further details on statistics. Schemes created with BioRender.com.

As expected from its ability to generate systemic inflammation^39^, we observed that an intraperitoneal (i.p.) injection of a low dose of LPS (0.02 mg kg^-1^) generated a significantly slow and mild fever response over hours with body temperatures typically not exceeding 38 °C (**Figures 1C, D, F, and S1C**). Furthermore, LPS-injected mice moved significantly less and had a slower walking speed than saline-injected animals (**Figures 1G, I, and S1D**). However, LPS-treated animals did not show an increase in stasis (periods >10 s without locomotion) compared with the controls (**Figure 1J and L**). The total amount of time mice interacted with food and the length of the interaction bouts (a proxy for food consumption or interest in food) during the four-hour post-injection period were not affected, but LPS-injected mice did interact with food at a lower rate (**Figures 1M, O, and S1E and F**).

As previously shown^33^, PGE2 (4 nmol i.c.v.) generated significantly high fever with body temperatures above 39 °C (**Figures 1C, E, and F, S1C**). In addition to its pyrogenic effect, PGE2 significantly reduced the walking speed in mice, but not the total distance traveled 90 minutes post-injection (**Figures 1H, I, and S1D**). Animals injected with PGE2 displayed significantly longer events of stasis (**Figure 1K and L**) and had a marked reduction in the overall time spent interacting with food, but not in the length of the interaction bouts (**Figures 1N, O, and S1E and F**). Altogether, we observed that PGE2 i.c.v. recapitulated many physiological and behavioral features observed during LPS-triggered sickness^1,2^ and natural infections^4,5,42^, including fever, slower locomotion, quiescence, and reduced eating. Crucially, these effects can be detected minutes after injections and are consistent with the role of PGE2 as one of the central effectors of LPS-triggered sickness^28^ and the chosen route of administration.

**Figure S1:**
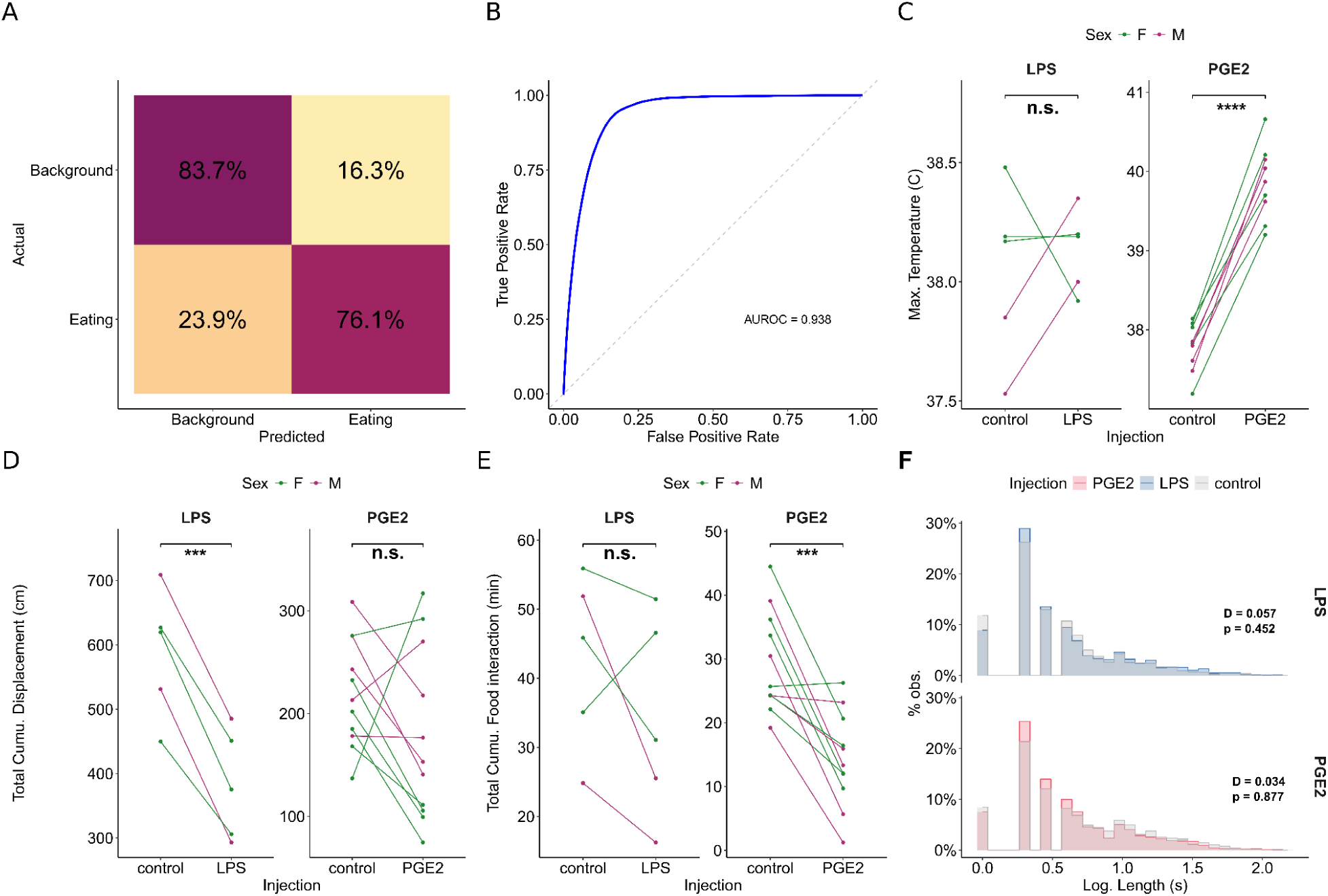
A central injection of PGE2 generates a robust sickness response in mice. **(A)** Confusion matrix for the deep learning-based food interaction detection model. **(B)** Receiver operating characteristic (ROC) curve for the food interaction classifier with area under the curve (AUROC) value. **(C-E)** Paired analysis of maximum body temperature, total cumulative movement, and cumulative food interaction time for PGE2 and LPS treatments versus controls (paired t-tests). Animals with missing paired values were dropped from the analysis (three mice in total). **(F)** Histograms showing the distribution of food interaction episode durations for PGE2 and LPS experiments compared to controls (Kolmogorov-Smirnov tests). See **Table 1** for further details on statistics.

### PGE2 activates the central autonomic network

Although LPS i.p. and PGE2 i.c.v. can both induce a comprehensive sickness state in mice (**Figures 1 and S1**), their mechanisms of action vary greatly. Peripheral inflammation caused by LPS is conveyed from the body to the brain partially through the peripheral nervous system and partially through the release of a complex mix of immune-borne humoral components into the bloodstream^9,12^. In contrast, the acute effect of PGE2 i.c.v. arguably bypasses any peripheral mechanisms and controls different aspects of sickness through direct interactions with central neurons expressing specific receptors for PGE2^8^. Given these differences, we evaluated to what extent the neuronal activation map triggered by LPS i.p. can be recapitulated by centrally-infused PGE2.

Multiple independent studies have evaluated the expression of the *Fos* gene, an early marker of neuronal activation^43^, across the mouse brain following i.p. administration of LPS. These studies show a remarkably consistent pattern of brain activation by LPS (**Figure S2A**)^1–3,13,14^. This pattern includes several hypothalamic nuclei, known for their roles in thermoregulation, stress, and energy homeostasis^35,45,46^, such as the preoptic area (POA), the paraventricular hypothalamus (PVH), and the arcuate nucleus (Arc). Additionally, in the brainstem, two regions involved in processing ascending spinal and visceral information^46,47^, such as the parabrachial nucleus (PBN) and the nucleus of the solitary tract (NTS), were also reported to respond to LPS in these studies. Key structures in the regulation of emotional states, consummatory behaviors, and interoceptive information integration^48–50^, including the central amygdala (CeA) and, to a lesser extent, the insular cortex (IC)^51^, also showed evidence of activation.

To compare with these LPS-induced activity maps, we quantified Fos expression 1.5 hours after delivery of PGE2 (4 nmol i.c.v.) in intact mouse brains utilizing the iDISCO approach^52,53^. The resulting whole-mount Fos expression patterns were registered to the CCFv3 Allen Mouse Brain Atlas, and expression densities were compared with vehicle-injected controls (Figure 2A, **Video S1, and Table S2**). Parts of the left IC showed significant activation after PGE2 injection (**Figures 2B and S2B**). In agreement with previous observations^1^, we also detected a significant reduction of Fos signal in specific thalamic nuclei such as the ventral medial nucleus (VMN) and the magnocellular part of the subparafascicular nucleus (PSPFm) (**Figures 2C and S2C**). Furthermore, hypothalamic nuclei rich in PGE2 receptors^33,54^, such as POA, PVH, and Arc, were found to be bilaterally activated by PGE2 (**Figures 2D and S2D**). In the midbrain, we noticed a significant increase in Fos+ cell densities within the lateral periaqueductal gray (PAG), an area associated with pain processing upon inflammation^55^ (**Figure 2E**). Additionally, brainstem regions that express ER receptors and are known to bind PGE2^32,56^, including regions of PBN, NTS, and neighboring areas, showed evidence of activation by PGE2 (**Figures 2F and S2F**). Overall, we found that PGE2 i.c.v. can activate many of the brain regions activated by LPS i.p. Furthermore, as observed during LPS-triggered sickness, the PGE2 activation map broadly encompassed the central autonomic network (CAN), a group of brain areas pivotal in conveying visceral sensory inputs into conscious interoceptive perceptions^48,49,57^.

**Figure 2:**
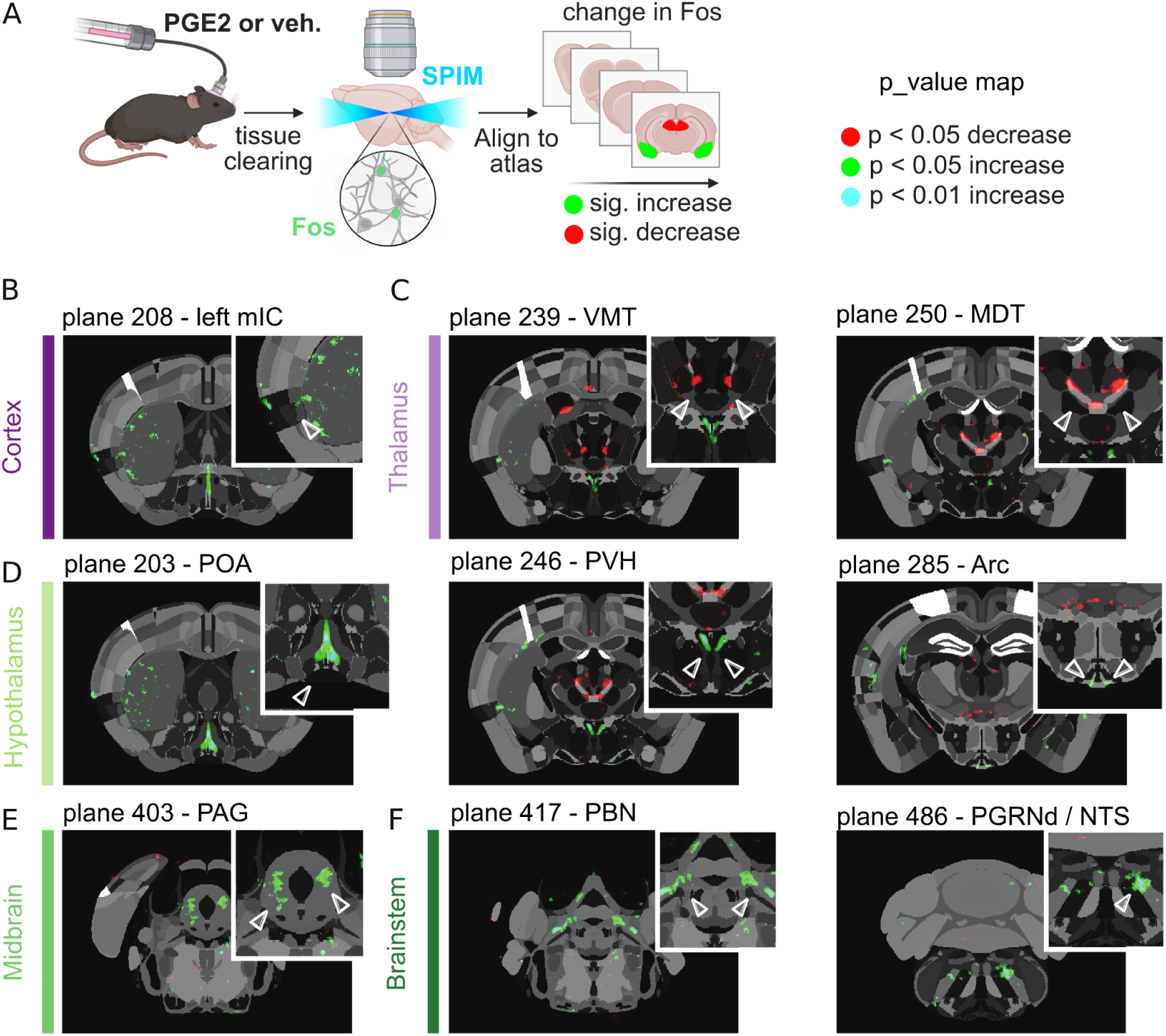
PGE2 activates the central autonomic network. **(A)** Schematic representation of the experimental approach. **(B-F)** p-value map of the difference between Fos staining signal density from vehicle-injected brains vs. PGE2-injected brains across selected brain planes from the Allen Brain Atlas. Different brain regions of interest are highlighted, including the cortex (B), thalamus (C), hypothalamus (D), midbrain (E), and brainstem (F). mIC: medial insular cortex; VMT: ventromedial nucleus of the thalamus; MDT: mediodorsal nucleus of the thalamus; POA: preoptic area; PVH: paraventricular hypothalamus; ARC: arcuate nucleus; PAG: periaqueductal gray; PBN: Paraventricular hypothalamic nucleus; PGRNd/NTS: Paragigantocellular reticular nucleus, dorsal part. Scheme created with BioRender.com.

**Figure S2:**
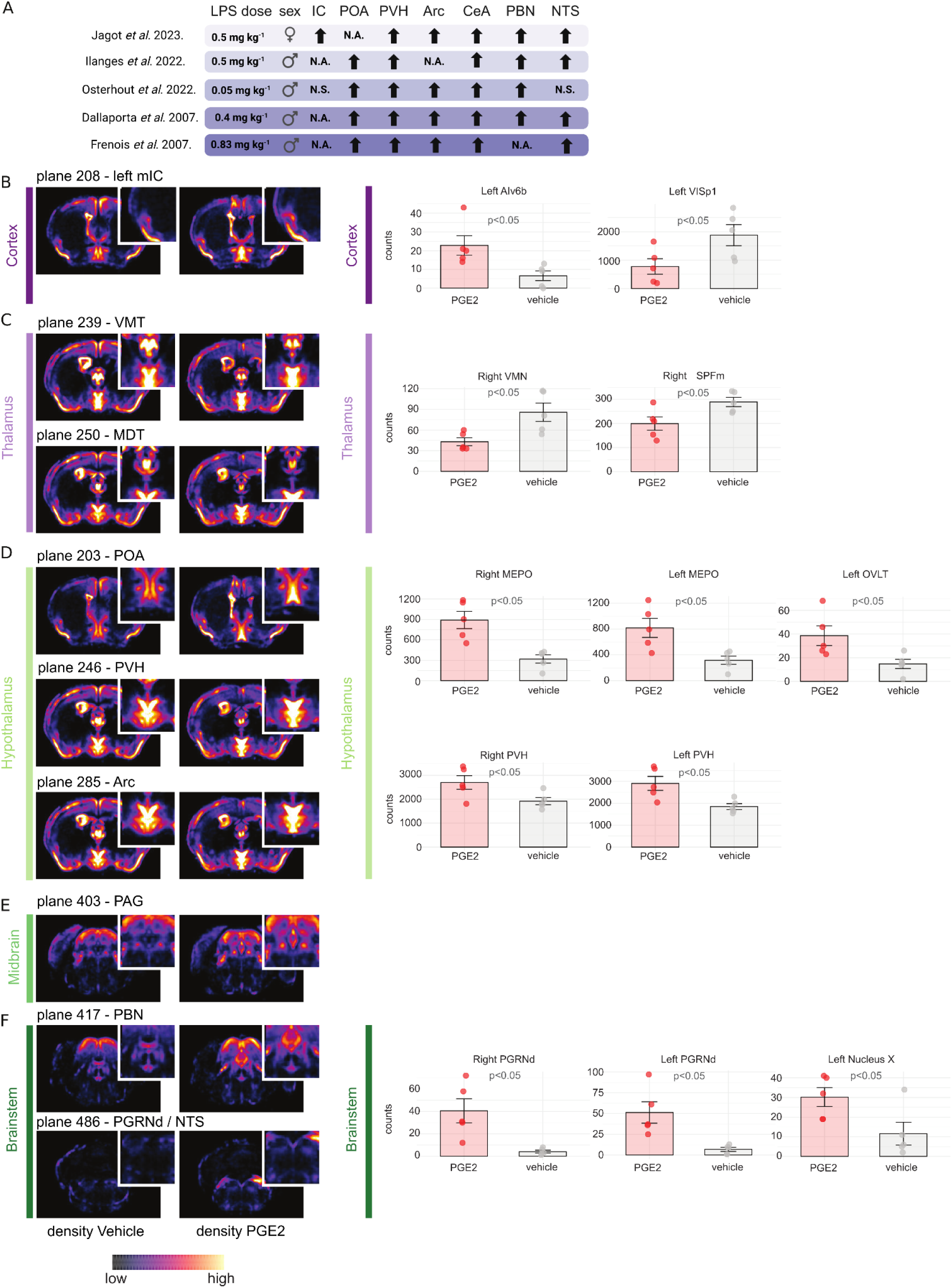
PGE2 activates the central autonomic network. **(A)** Schematic summary of selected reports on the effects of LPS injection on Fos levels across the mouse brain. Upward pointing arrow: significant increase in Fos levels. N.S: not significant. N.A: Not available. **(B-F)** Left Panel: Fos staining signal density across selected brain planes from the Allen Brain Atlas for vehicle-injected brains and PGE2-injected brains (same planes used for Fig. 2 p-value maps). Different brain regions of interest are highlighted, including the cortex (B), thalamus (C), hypothalamus (D), midbrain (E), and brainstem (F). Right panel: Fos-positive nuclei counts across the respective brain areas in the left and right hemispheres of PGE2 and vehicle-injected mice. The brain regions of interest span the cortex (B), thalamus (C), hypothalamus (D), and brainstem (F). See **Table S2** for further details on statistics. mIC: medial insular cortex; Alv6b: agranular insular area, ventral part, layer 6b; VISp1: primary visual area; VMT: ventromedial nucleus of the thalamus; MDT: mediodorsal nucleus of the thalamus; SPFm: subparafascicular nucleus, magnocellular part; POA: preoptic area; PVH: paraventricular hypothalamus; ARC: arcuate nucleus; MEPO: median preoptic nucleus; OVLT: organum vasculosum of the lamina terminalis; PAG: periaqueductal gray; PBN: paraventricular hypothalamic nucleus; PGRNd/NTS: paragigantocellular reticular nucleus, dorsal part. Table scheme created with BioRender.com.

### Single-unit analysis of the neuronal correlates of sickness in head-fixed mice

In humans, sickness is perceived as a discrete, low-valence state, distinguishable from other states^58^. This perception is accompanied by the activation of brain regions involved in processing interoceptive information, suggesting that sickness generates a distinct body state representation^10^. Our results show that PGE2 i.c.v. triggers a fast sickness response in mice (**Figures 1 and S1**), and the PGE2-associated physiological and behavioral changes are accompanied by activation of a large portion of the CAN (**Figures 2 and S2**). These two characteristics make an infusion of PGE2 i.c.v. a powerful tool to investigate the neuronal dynamics underlying potential changes in body state during sickness in awake mice. Among the brain areas proposed to participate in the representation of bodily states, including immune status, are the insular cortex (IC) and the primary somatosensory cortex (SSp)^15,48,59^. However, we currently have limited information about neuronal activity in these cortical areas during inflammation at the level of single-neuron responses.

To address this gap, we developed a methodology to collect single-neuron electrophysiological data in awake mice while infusing PGE2 i.c.v. (**Figure 3A, Methods**). We obtained high-density neuronal recordings of the IC and SSp using Neuropixel probes^60^ while monitoring body temperature and behavior in head-fixed mice (**Figure 3B, C, and D**). Our single-unit analysis revealed that PGE2 does not induce gross changes in neuronal activity patterns in the IC and SSp. However, we noted a slight and general decrease in the average firing frequency (FF) compared with vehicle infusion (**Figure S3A and B**).

**Figure 3:**
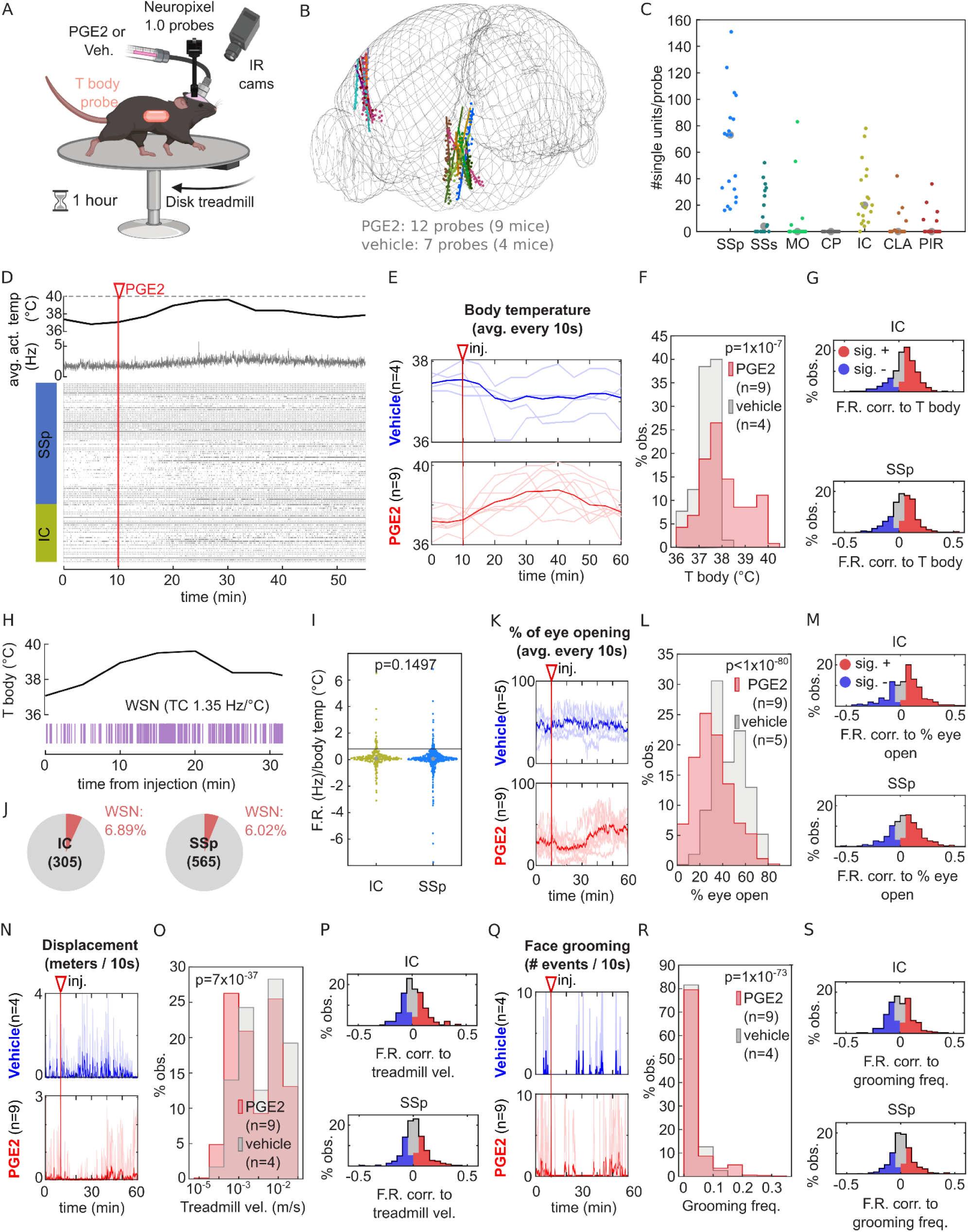
Single-unit analysis of the neuronal correlates of sickness in head-fixed mice. **(A)** Schematic representation of the experimental design. **(B)** Reconstruction of the electrode tracks from recorded mice aligned to the brain atlas. **(C)** Number of single units recorded per brain area. Each point represents data from one probe insertion. SSp: principal somatosensory cortex; SSs: secondary somatosensory cortex; MO: primary motor cortex; CP: caudate putamen; IC: insular cortex; CLA: claustrum; PIR: piriform cortex. **(D)** Representative dataset. Top panel: body temperature trace. Middle panel: Average population firing rate (F.R.). Bottom panel: Raster plot of simultaneously recorded single units across SSp (blue) and IC (yellow) with one probe. Red line: time of injection of PGE2. **(E)** Body temperature recorded across the experiment for vehicle (top) and PGE2-injected mice (bottom). The values plotted were averaged across 10s time bins. Light color traces correspond to individual mice, and solid color traces are the average across mice. The highlighted vertical line points out the injection time. **(F)** Histograms of pooled body temperatures recorded in response to the injections, excluding the 10 minutes pre-injection and 2 minutes during injection, from 9 PGE2-injected and 4 vehicle-injected mice (red and gray, respectively). Statistics: 2-sample Kolmogorov-Smirnov test. **(G)** Histograms of the correlation coefficients between recorded single units’ firing rate dynamics and body temperature across the experiment for IC (top) and SSp (bottom) from PGE2-injected mice. Red areas in the histogram represent significantly positive correlations, and blue areas significantly negative correlations (Pearson’s correlation coefficient p < 0.05). **(H)** Top: Body temperature trace. Bottom: Firing pattern of a representative warm-sensitive neuron (WSN). TC: temperature coefficient (change in F.R / °C). **(I)** Firing rate change per degree of body temperature (TC) calculated for each recorded single unit from IC (yellow, 305 cells) and SSp (blue, 565 cells). Data from 9 PGE2-injected mice are plotted. Dotted lines represent the TC limit for WSN (TC = 0.8 Hz/°C). Median TC for IC = 0.092; median TC for SSp = 0.063. Statistics: Mann-Whitney test. **(J)** Pie charts showing the percentage of WSNs detected in IC (left) and SSp (right). The data displayed in I) were used for these plots. **(K-M)** Same as (E-G) but for % of eye opening. **(N-P)** Same as (E-G) but for treadmill velocity. **(Q-S)** Same as (E-G) but for grooming frequency. See **Table 1** for further details on statistics.

Fever is arguably the most well-known indicator of an infectious disease, and is routinely used to monitor illness progression^7^. Therefore, as a starting point, we used the body temperature fluctuations triggered by PGE2 as a proxy for sickness. As observed in freely-moving mice, PGE2 i.c.v. triggers a rapid fever response in mice with temperatures reaching 40 °C (**Figure 3E and F**). We observed that IC and SSp had comparable proportions of neurons whose activity was significantly correlated (positively and negatively) with body temperature (**Figure 3G, Table S1**). Warm-sensitive neurons (WSN) are characterized by a steep increase in their firing rate in response to temperature changes and are proposed to detect brain temperature^61–63^. The proportions of these WSNs were similar between the IC and SSp (**Figure 3H and I**), and lower than the main central thermoregulatory region in the hypothalamus^61^. Furthermore, neurons in both regions exhibited an average temperature coefficient close to 0 Hz/°C (0.096 and 0.063 Hz/°C, respectively) (**Figure 3J, Table S1**). Altogether, these data suggest that potential changes in brain activity after PGE2 injection are not solely driven by a passive increase in FF due to the temperature sensitivity (positive Q_10_) of most ion channels involved in synaptic transmission^64^.

Next, we focused on the behavioral features associated with PGE2-triggered sickness. We observed that PGE2-injected mice squinted their eyes (**Figure 3K and L**), a facial expression linked to physical pain in laboratory animals^65^ and humans^66^. Furthermore, PGE2-treated mice walked significantly slower (**Figure 3N and O**) and exhibited increased face grooming, compared to vehicle-infused controls (**Figure 3Q and R**). As with fever, IC and SSp had comparable proportions of individual neurons whose activity correlated (both positively and negatively) with these three behaviors (**Figure 3M, P, and S, Table S1**).

Altogether, we found that PGE2 induces a rapid and comprehensive sickness response in head-fixed mice, similar to what is observed in freely moving animals. Furthermore, we detected heterogeneous changes in neuronal activity in the SSp and IC, with groups of neurons showing both positive and negative correlations with sickness symptoms.

**Figure S3:**
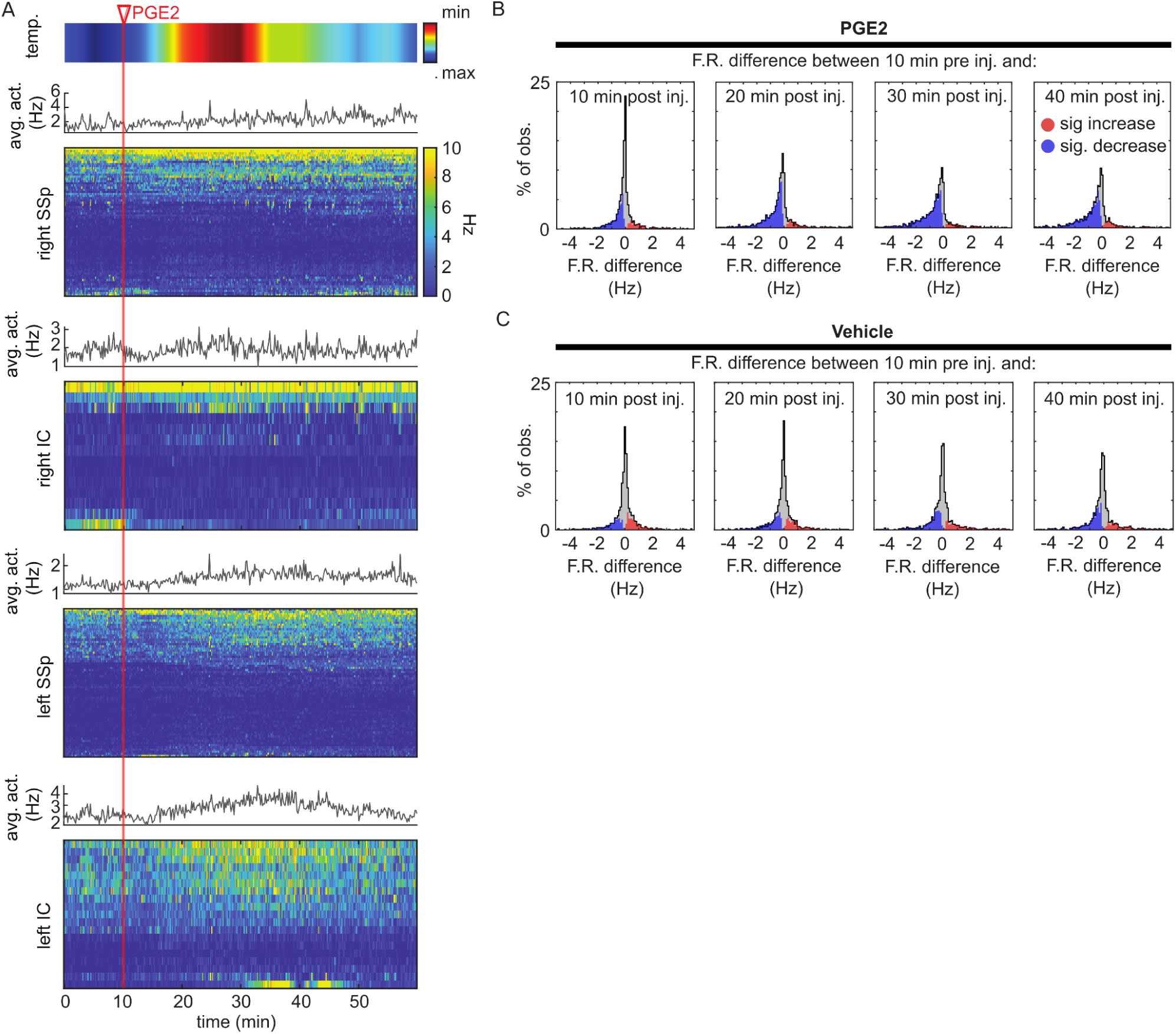
Single-unit analysis of the neuronal correlates of sickness in head-fixed mice. **(A)** Representative dataset of the firing rates across the experiment from simultaneous bilateral single-unit recordings in the left and right SSp and IC. Firing rates were computed over 10s time bins. Each firing rate matrix shows the corresponding averaged population firing rate on top. The top panel shows the body temperature response to PGE2 injection. The highlighted vertical line shows the injection time. **(B)** Histograms of the firing rate differences between the 10 minutes pre-injection and time windows consisting of 10, 20, 30, or 40 minutes post-injection, across the recorded single-units from 9 PGE2 injected mice. Red areas in the histogram represent significantly higher firing rates post-injection, and blue areas significantly lower firing rates post-injection (2-sample Kolmogorov-Smirnov tests between the firing rates evaluated every second, across 10 min before and after injection, p < 0.05). **(C)** Same as (B), but for 4 vehicle-injected mice. See **Table 1** for further details on statistics.

### A sickness-associated state across neocortical regions

The study of the firing properties of individual neurons has greatly contributed to our understanding of brain function. However, there is a growing consensus that low-dimensional patterns of population neuronal activity may play a crucial role in complex brain functions, such as encoding sensory stimuli, motor tasks, and internal states^67–71^. To test whether sickness-related variables could be identified in low-dimensional representations of our neuronal recordings, we used CEBRA^72^. CEBRA is a supervised nonlinear dimensionality reduction method that enables the identification of consistent low-dimensional representations of neuronal activity (known as embeddings) across data collected from different animals, using behavioral information as guidance. As a result, neuronal population activity that substantially explains observed behavioral patterns produces well-defined structures in the 3D embeddings generated by CEBRA.

Using the multi-session CEBRA-behavior algorithm, we mapped the neuronal activity from all recorded regions (All), the SSp, or the IC to CEBRA embeddings generated considering body temperature alone, sickness behaviors (eye-opening, walking speed, and grooming), or a combination of all factors. We then trained a support vector machine (SVM) classifier to determine whether the embedded neuronal activity could be used to distinguish between PGE2 and vehicle conditions.

We observed well-defined structures in the 3D CEBRA embeddings across all three categories studied, both in the full dataset and in subsets containing only units from the IC and SSp (**Figure 4A, S4A and B**). The binary SVM classifiers decoded the injection type (PGE2 or vehicle) significantly above chance (scrambled labels on embedded data) for all comparisons, with fractions of correctly classified time bins (hit rate) ranging from 0.76 to 0.92 (**Figure 4B, Table S1**). Interestingly, decoders trained on IC data outperformed those trained on SSp data, particularly when body temperature or all factors were used as auxiliary variables (**Figure 4B**). These results suggest that PGE2 induces discernible changes in neuronal population dynamics across all recorded regions, a characteristic associated with bodily states^73^. We found that low-dimensional patterns of neuronal activity may underlie the sickness features exhibited by the animals, supporting the existence of a sick state. Furthermore, among the studied regions, the IC harbors a better representation of body temperature and its combination with sickness-related behaviors than the SSp.

**Figure 4:**
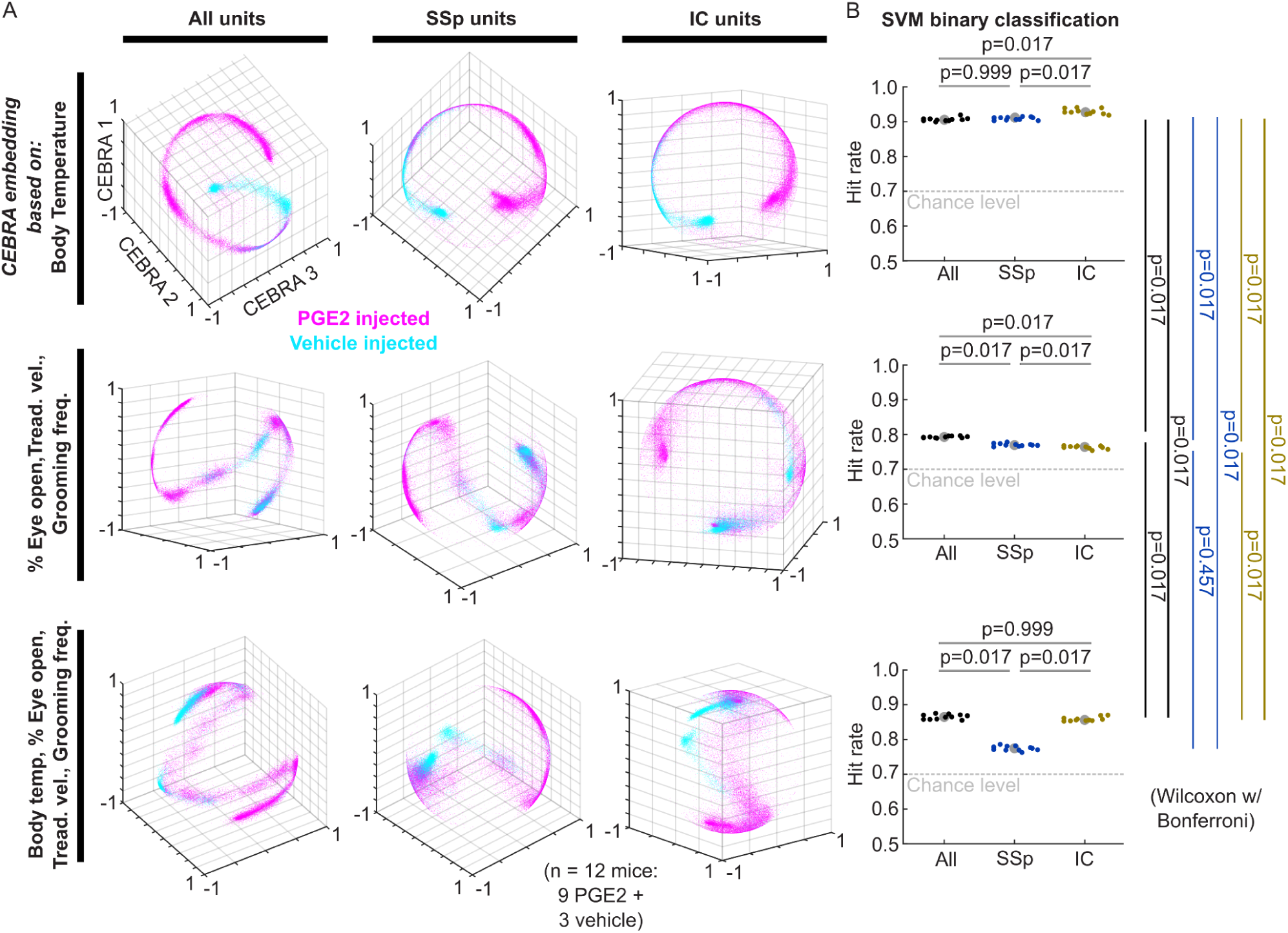
A sickness-associated state across neocortical regions. **(A)** 3D embeddings of the firing rates recorded in 1s time bins (plotted points) across 48 minutes post-injection of either vehicle (cyan, n = 3 mice, 5 probes) or PGE2 (magenta, n = 9 mice, 12 probes) obtained through multi-session CEBRA-behavior supervised dimensionality reduction (see methods). Embeddings for data from all units simultaneously recorded from each probe across brain regions (left) or specifically from SSp (center) or IC (right) are plotted. Multi-session CEBRA-behavior embeddings were obtained using either body temperature (top), the three behavioral readouts studied (middle), or all 4 readouts together (bottom), as accessory variables. **(B)** Hit rates for predicting the correct treatment labels (vehicle or PGE2) obtained through cross-validated binary classification of the corresponding embedded points in (A) using a support vector machine classifier (SVM). The chance level hit rate (Chl) obtained by scrambling the treatment labels is plotted in gray dotted lines. Statistics: Wilcoxon’s signed rank tests with Bonferroni correction for multiple comparisons.

Multiple studies have reported a certain level of lateralization in engaged neocortical regions, including the IC, during systemic inflammation in both humans^10,17,74^ and mice^51^. Consistently, our Fos-iDISCO map showed an enrichment of Fos signal, particularly in the left IC after PGE2 i.c.v. (**Figure 2B and S2B**). Based on this evidence, we explored the respective contributions of the left and right hemispheres of the SSp and IC to sickness representation. Interestingly, decoders trained on data from the left hemisphere outperformed those trained on the right when using body temperature as an accessory variable across both SSp and IC regions. A similar pattern was observed in the IC when combining body temperature with sickness behaviors. Conversely, sickness behaviors alone were better decoded from embeddings obtained from the right hemisphere in both SSp and IC (**Figure S4C and D**). These results suggest a degree of lateralization in sickness representation within the IC and SSp, with different aspects of sickness preferentially represented in either the right or left hemisphere.

**Figure S4:**
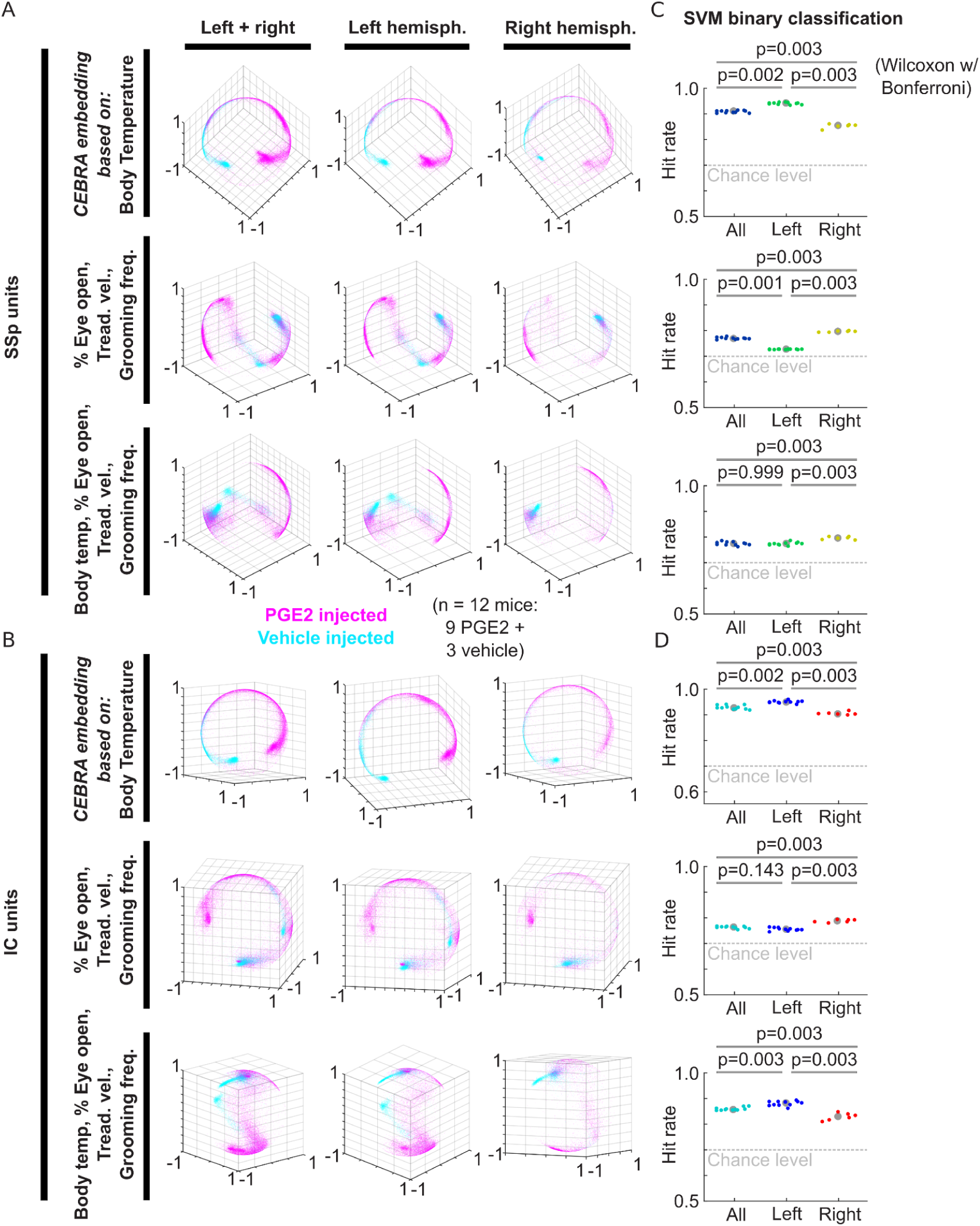
A sickness-associated state across neocortical regions. **(A)** 3D embeddings of the firing rates recorded in 1s time bins (plotted points) across 48 minutes post-injection of either vehicle (cyan, n = 3 mice, 5 probes) or PGE2 (magenta, n = 9 mice, 12 probes) obtained through multi-session CEBRA-behavior supervised dimensionality reduction (see methods). Embeddings for data from units located at SSp simultaneously recorded from each probe across brain regions (Left + right) or specifically from probes implanted in the left or right hemispheres are plotted. Multi-session CEBRA-behavior embeddings were obtained using either body temperature (top), the three behavioral readouts studied (middle), or all 4 readouts together (bottom), as accessory variables. **(B)** Same as (A) but for units located at IC exclusively. **(C)** Hit rates for predicting the correct treatment labels (vehicle or PGE2) obtained through cross-validated binary classification of the corresponding embedded points in (A) using a support vector machine classifier (SVM). The chance level hit rate (Chl) obtained by scrambling the treatment labels is plotted in gray dotted lines. Statistics: Wilcoxon’s signed rank tests with Bonferroni correction for multiple comparisons. **(D)** Same as (C) but for units located at IC exclusively.

### A divergent representation of sickness behaviors between IC and SSp

CEBRA enables the integration of neuronal recordings from different individuals by transforming measured neuronal activity into a shared embedding across experimental animals. However, to rule out a potential influence from biases in the consistency of CEBRA-generated embeddings (**Figure S5A**), we explored an alternative method to validate our findings.

To further assess how neuronal population dynamics in IC and SSp account for changes in body temperature and behavior following PGE2 infusion, we performed a decoding analysis using generalized linear models (GLM) on the direct measurements of neuronal activity during sickness. We found that models trained on the data from IC performed significantly better (overall smaller prediction errors) at predicting body temperature fluctuations than those trained on the SSp data (**Figure 5A**). Furthermore, decoders trained on brain activity from areas in the left hemisphere performed better at predicting body temperature than those trained on data collected from the right hemisphere (**Figure 5B and S5B**). Within the left hemisphere, population dynamics of the left IC allowed a better decoding performance than those trained on data from the left SSp (**Figure 5C**). In line with the CEBRA findings, the GLM analysis suggests that information from ventrally and left-located brain areas yields more accurate body temperature predictions. Specifically, the left IC demonstrated a more accurate representation of body temperature following PGE2 injection.

**Figure 5:**
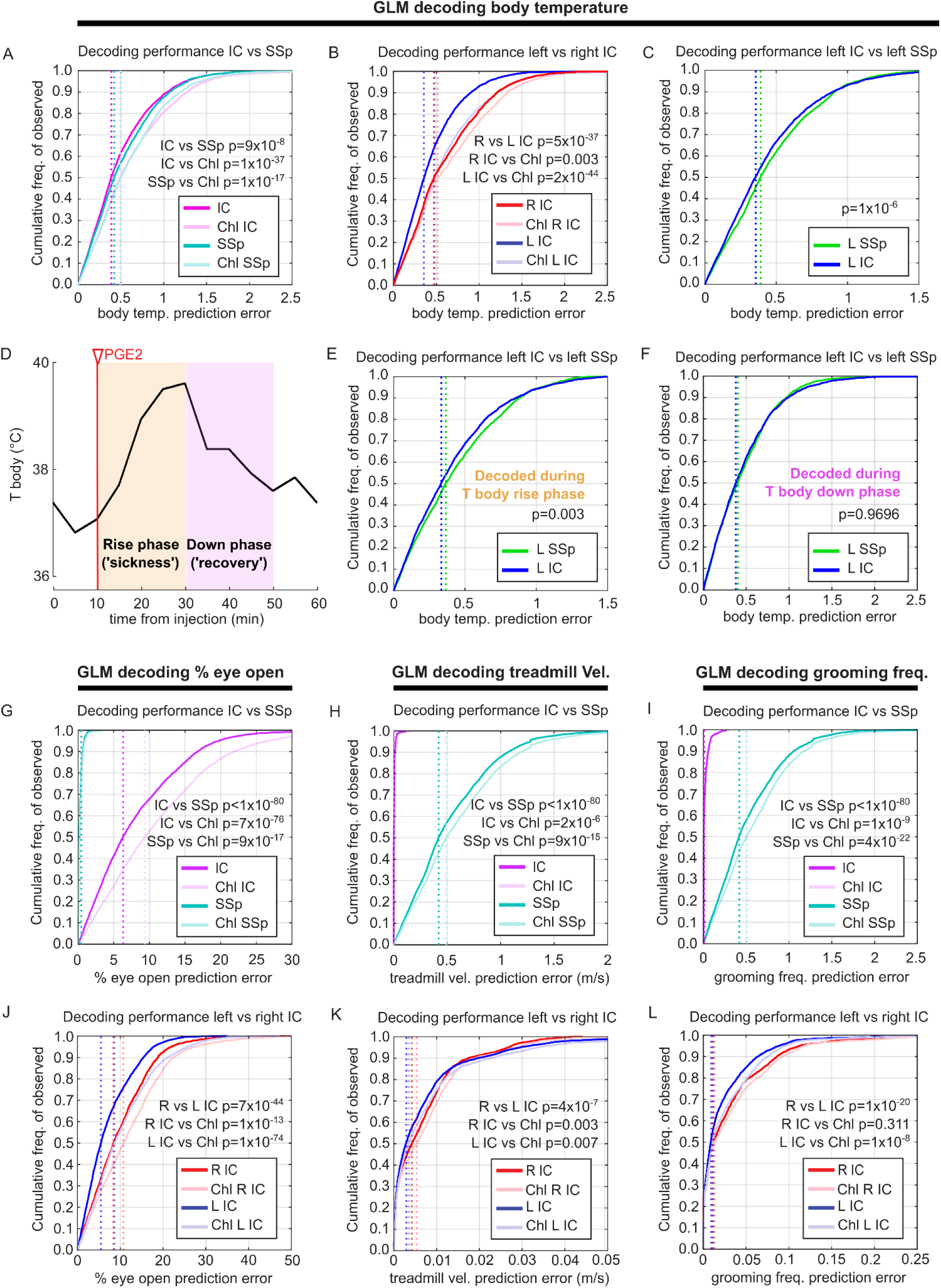
A divergent representation of sickness behaviors between the IC and SSp. **(A)** Cumulative distribution plots of the decoding errors (°C, T body - predicted) obtained using GLM’s on the simultaneously recorded IC populations (magenta) and corresponding SSp populations of equal size (cyan), across the datasets from 9 PGE2-injected mice. The chance level distributions (Chl) obtained by scrambling the temperature traces are plotted in light magenta and light cyan, respectively. Vertical dotted lines correspond to each distribution’s median. Statistics: 2-sample Kolmogorov-Smirnov tests with Bonferroni correction for multiple comparisons. **(B)** Same as in (A) but comparing the IC decoding error distributions from probes implanted in the right hemisphere (red, n = 4) versus the left hemisphere (blue, n = 8). Respective chance levels are in light red and blue. Same statistics as J). **(C)** Same as in (A) but for probes implanted in the left hemisphere (n = 8), comparing the decoding error distributions obtained from the left IC (blue) and the left SSp (green). Same statistics as (A). **(D)** Representative body temperature response to PGE2 injection, demarcating the fever rise phase and down phase. **(E, F)** Same as in (C), but decoding IC population dynamics during the temperature rise phase **(E)** and down phase **(F)**. Same statistics as (A). **(G-I)** Cumulative distribution plots of the decoding errors for eye-opening percentage **(G)**, treadmill velocity **(H),** and grooming frequency (mean #events/sec. in 5 min, **I**) obtained using GLM’s on the simultaneously recorded IC populations (magenta) and corresponding SSp populations of equal size (cyan) across the datasets from 9 PGE2-injected mice. The chance level distributions obtained by scrambling each behavioral readout are plotted in light red and light blue, respectively. Vertical dotted lines correspond to each distribution’s median. Statistics: 2-sample Kolmogorov-Smirnov tests with Bonferroni correction for multiple comparisons. **(J-L)** Same as in (G-I) but comparing the IC decoding error distributions from probes implanted in the right hemisphere (red, n = 4) versus the left hemisphere (blue, n = 8). Respective chance levels are in light red and blue, same statistics as (G). See **Table 1** for further details on statistics.

Next, we separately analyzed the population dynamics during the upward and the downward phases of the fever response, as they likely represent different body states: sickness and recovery, respectively (**Figure 5D**). Interestingly, the better decoding performance of body temperature information from the left IC over the left SSp was only detected during the upward phase of fever (**Figure 5E and F**). This suggests that the information carried by the left IC during the sickness phase is the primary factor driving the better decoding performance of body temperature fluctuations observed in the left IC compared to the left SSp. On the other hand, the difference in decoding performance between right and left-located brain areas was preserved in both phases between IC and SSp (**Figure S5D to I**).

We then extended our GLM decoding analysis to predict individual sickness behaviors from neuronal population dynamics recorded in the IC and SSp. Decoders trained on the data from IC and SSp predicted each of the three behaviors better than chance. However, eye squinting was more accurately decoded using the data from SSp while walking speed and grooming were better predicted from IC (**Figure 5G, H, and I**). Similar to the pattern observed for fever, decoding performance was better from data obtained from the left hemisphere in both IC and SSp (**Figures 5J, K, and L and S5J, K, and L**).

**Figure S5:**
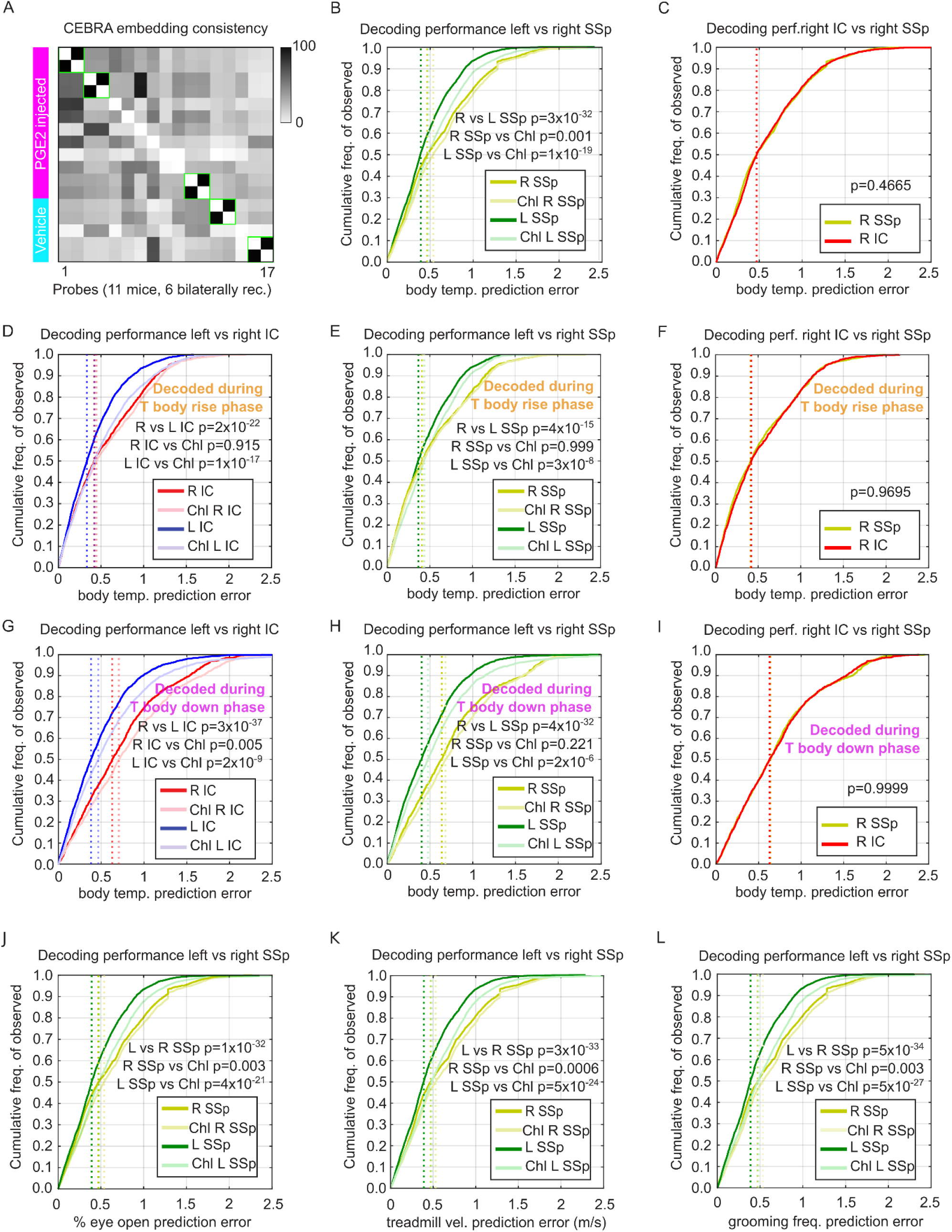
A divergent representation of sickness behaviors between the IC and SSp. **(A)** Representative CEBRA-behavior embedding consistency matrix across datasets (simultaneously recorded unit firing rates across probes) for data from both brain hemispheres (All) and using body temperature as an accessory variable for embedding. Green squares highlight consistent data from 5 bilaterally recorded mice. **(B)** Cumulative distribution plots of the decoding errors for T body obtained by GLM’s on the simultaneously recorded SSp populations from the right hemisphere (yellow, n = 4) versus the left hemisphere (green, n = 8). Respective chance levels are in light red and blue. Vertical dotted lines correspond to each distribution’s median. Statistics: 2-sample Kolmogorov-Smirnov tests with Bonferroni correction for multiple comparisons. **(C)** Same as in B) but for probes implanted in the right hemisphere (n = 4), comparing the decoding error distributions obtained from the right IC (red) and the right SSp (yellow). Same statistics as (B). **(D)** Same as in (B) but comparing the IC decoding error distributions obtained during the body temperature rise phase from probes implanted in the right hemisphere (red, n = 4) versus the left hemisphere (blue, n = 8). Respective chance levels are in light red and blue. Same statistics as (B). **(E)** Same as in (B) but comparing the SSp decoding error distributions during the body temperature rise phase from probes implanted in the right hemisphere (yellow, n = 4) versus the left hemisphere (green, n = 8). Same statistics as (B). **(F)** Same as in B) but for probes implanted in the right hemisphere (n = 4), comparing the decoding error distributions obtained during the body temperature rise phase from the right IC (red) and the right SSp (yellow). Same statistics as (B). **(G-I)** is the same as (D), (E), and (F), respectively, but for the body temperature down phase. **(j-L)** Cumulative distribution plots of the decoding errors for eye opening percentage **(J)**, treadmill velocity **(K),** and grooming frequency **(L)** obtained using GLM’s on the simultaneously recorded SSp populations from the right hemisphere (yellow, n = 4) versus the left hemisphere (green, n = 8). Respective chance levels are in light red and blue. Vertical dotted lines correspond to each distribution’s median. Statistics: 2-sample Kolmogorov-Smirnov tests with Bonferroni correction for multiple comparisons. See **Table 1** for further details on statistics.

### The IC encodes a unified readout of sickness, outperforming fever alone

So far, our GLM approach has examined different behavioral and physiological traits of PGE2-triggered sickness separately. However, as highlighted by our CEBRA analysis, all physiological and behavioral symptoms contribute to the overall sick state. To evaluate their joint contribution, we defined a unified readout of sickness, named the sickness index (SI). To calculate the SI, we first normalized the following measures: body temperature, the inverse of eye squinting, the inverse of walking velocity (both inversely proportional to PGE2-induced sickness), and grooming. Given no a priori reason to differentially weight these symptoms in the sick state, we normalized their sum to define the SI. The SI generally tracked body temperature profiles, although its temporal dynamics differed, with the SI plateauing earlier on average (**Figures 6A and 3E**). Moreover, PGE2-treated mice exhibited a significantly higher SI than vehicle-infused controls (**Figure 6B**). These findings prompted us to use SI as a readout for the sick state.

**Figure 6:**
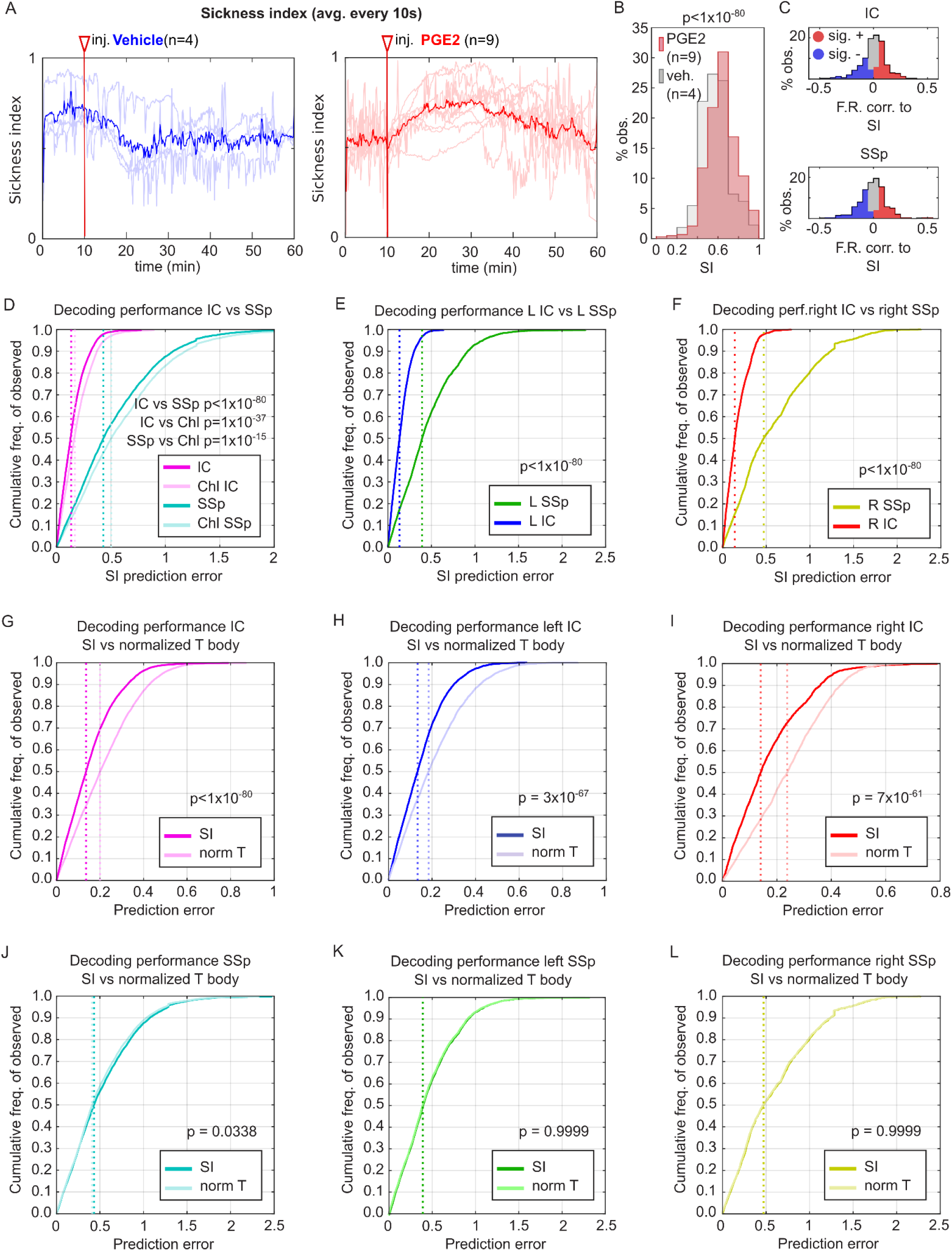
The IC encodes a unified readout of sickness, outperforming fever alone. **(A)** Sickness index recorded across the experiment for vehicle (left) and PGE2-injected mice (right). The values plotted were averaged across 10s time bins. Light color traces correspond to individual mice, and solid color traces are the average across mice. The highlighted vertical line points out the injection time. **(B)** Histograms of sickness index recorded in response to the injections, excluding the 10 minutes pre-injection and 2 minutes during injection, from 9 PGE2-injected and 4 vehicle-injected mice (red and gray, respectively). Statistics: 2-sample Kolmogorov-Smirnov test. **(C)** Correlation coefficient histograms for single units’ firing rate dynamics and sickness index for IC (top) and SSp (bottom). Red areas in the histogram represent significantly positive correlations, and blue areas significantly negative correlations (Pearson’s correlation coefficient p < 0.05). **(D)** Cumulative distribution plots of the decoding errors (SI - predicted) obtained using GLM’s on the simultaneously recorded IC populations (magenta) and corresponding SSp populations of equal size (cyan), across the datasets from 9 PGE2-injected mice. The corresponding chance level distributions are plotted in light magenta and light cyan. Vertical dotted lines correspond to each distribution’s median. Statistics: 2-sample Kolmogorov-Smirnov tests with Bonferroni correction for multiple comparisons. **(E)** Same as in (D) but for probes implanted in the left hemisphere (n = 8), comparing the decoding error distributions obtained from the left IC (blue) and the left SSp (green). Same statistics as (D). **(F)** Same as in (E) but for probes implanted in the right hemisphere (n = 4), comparing the decoding error distributions obtained from the right IC (red) and the right SSp (yellow). **(G)** Comparison of the decoding error distributions for SI (magenta) and body temperature normalized to the same scale as SI (0 to 1 range, light magenta), obtained using GLM’s on the simultaneously recorded IC populations (magenta) across the datasets from 9 PGE2 injected mice. Vertical dotted lines correspond to each distribution’s median. Statistics: 2-sample Kolmogorov-Smirnov tests with Bonferroni correction for multiple comparisons **(H)** Same as in (G) but for probes implanted on the left hemisphere (n = 8). **(I)** Same as in (G) but for probes implanted in the right hemisphere (n = 4), **(J-L)** Same as (G-I), respectively, but for corresponding SSp populations of equal size to the IC populations studied.

Individual neuronal analysis indicated that IC and SSp had comparable proportions of neurons with activity positively or negatively correlated with SI (**Figure 6C, Table S1**). At the population level, we found that information about SI can be decoded better than chance level from all studied areas. However, we observed that models trained on IC data performed remarkably better than those trained on SSp data (**Figure 6D**). Interestingly, the superior SI encoding by the IC compared to SSp was evident in both hemispheres, with data from both the right and left IC yielding better SI predictions than their SSp counterparts (**Figure 6E and F**). Furthermore, left-hemisphere data consistently provided better decoding performance than right-hemisphere data for both IC and SSp recordings (**Figure S6A and B**).

Intrigued by the remarkable difference in SI decoding performance between IC and SSp, we next compared the performance of decoders trained on SI versus normalized body temperature alone. We found that the neuronal population activity of IC encoded SI better than the temperature alone (**Figure 6G**), and this was the case for both the left and right IC (**Figure 6H and I**). In contrast, only a modest improvement of decoding performance was achieved with the SSp data (**Figure 6J**), and this effect was not observed in either the left or right SSp alone (**Figure 6K and L**).

Altogether, in line with the proposed role of IC in representing convergent information about physiological states^75^, we found a more prominent encoding of sickness in IC than in SSp. Importantly, this preferential representation of sickness in IC is most clearly revealed when employing a unified measure of the sick state (represented here by the SI) rather than individual sickness readouts, such as fever.

**Figure S6:**
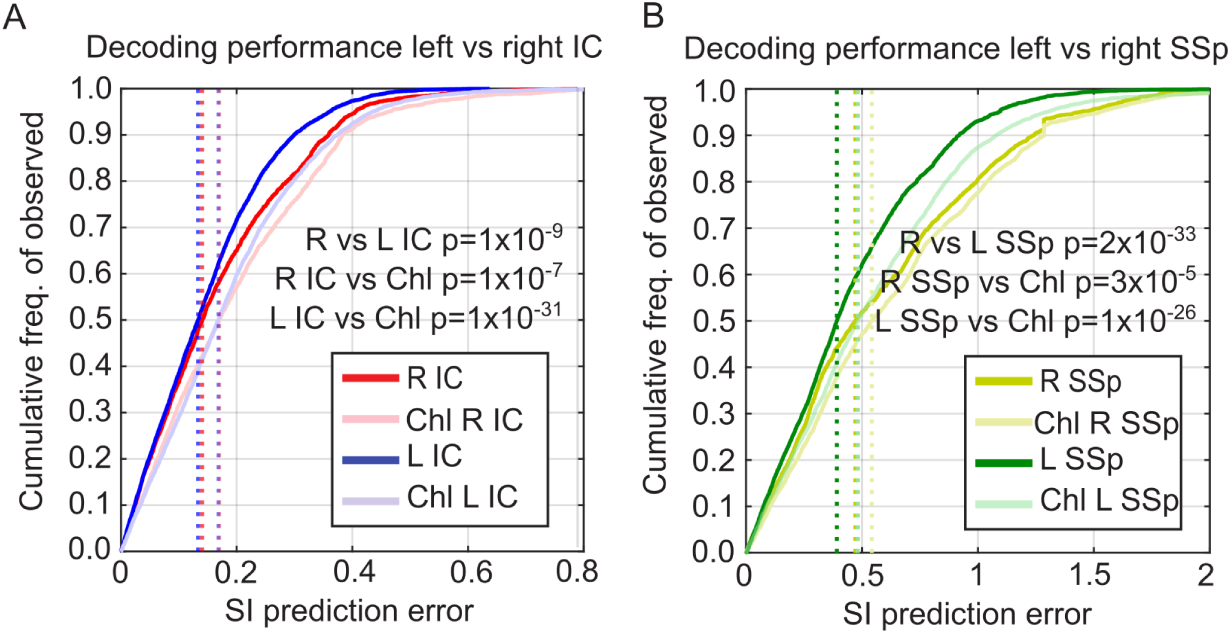
The IC encodes a unified readout of sickness, outperforming fever alone. **(A)** Cumulative distribution plots of the decoding errors for SI obtained using GLM on the simultaneously recorded IC populations from the right hemisphere (red, n = 4) versus the left hemisphere (blue, n = 8). Respective chance levels are in light red and blue. Vertical dotted lines correspond to each distribution’s median. Statistics: 2-sample Kolmogorov-Smirnov tests with Bonferroni correction for multiple comparisons. **(B)** Same as in (A) but for SSp.

## Discussion

Our study introduces a novel methodology to investigate real-time changes in body state associated with inflammation and its neural activity correlates in awake, behaving mice. We applied this approach to examine the neuronal correlates of sickness in the neocortex. Our behavioral analysis demonstrated that a single injection of PGE2 i.c.v. generates a robust sickness response in mice, which includes fever, slower locomotion, quiescence, anorexia/lack of motivation to eat, and eye squinting. All these physiological and behavioral features have been observed in a wide range of vertebrate species, including humans, during systemic inflammation and illness^4,5,7,42,65,66^. The strong effect of PGE2 is consistent with its role as one of the final immune mediators of sickness within the central nervous system (CNS)^8^. Indeed, during systemic inflammation, PGE2 production in the brain increases steeply, and the concentration of PGE2 in the cerebrospinal fluid is tightly correlated with sickness progression^27,28,76^.

In addition to its strong effect on behavior, we observed that PGE2 induces a distinct pattern of Fos expression in the brain that broadly aligns with the central autonomic network (CAN), a diverse group of brain regions involved in generating conscious interoceptive perceptions from visceral sensory inputs^48,49,57^. Furthermore, many brain areas enriched in Fos expression after PGE2 injection have consistently been reported to respond to systemic inflammation and infections in mice and humans^1–3,10,13,14,17^. Additionally, we found that PGE2 reduces Fos expression in diverse thalamic nuclei, an observation also seen during LPS-triggered peripheral inflammation^1^. Interestingly, among the nuclei showing evidence of decreased Fos signal upon PGE2 injection are important relay stations for ascending sensory inputs from the viscera to IC, such as the ventral medial nucleus (VMN) and, to a lesser extent, the ventral posteromedial parvicellular nucleus of the thalamus (VPMpc)^65,66^. The degree and implications of a potential reduction of activity in the thalamus during sickness will require further study.

Although PGE2 and LPS produce similar activation patterns in the brain, the mechanisms by which they impact brain activity differ. Systemic inflammation, as triggered by LPS, reaches the brain through the peripheral nervous system (PNS) and via the release of immune mediators into the bloodstream. Some of these mediators will ultimately trigger prostaglandin production in the brain, including PGE2^9,12,28^. Consistent with this dual communication route, resection of the vagus nerve, one of the main routes for ascending visceral information to the CNS, can reduce symptoms of sickness and lower activity in the CNS upon injection of specific doses of LPS i.p. ^12,76^. In contrast, PGE2 i.c.v. likely bypasses any peripheral mechanisms and acts directly on brain areas that express EP receptors. In agreement with this, our PGE2 activation map includes multiple areas reported to express EP receptors or bind PGE2 directly, such as POA, PVH, Arc, PBN, and NTS^32,33,54,56^.

A limitation of this study is that the injection of PGE2 i.c.v. is a reductionist approach, as it overlooks the contribution of many important humoral mediators and visceral sensory inputs present during infections. As such, it assumes that the effects of PGE2 result primarily from the activation of EP receptors directly in the brain, with little to no contribution from inputs originating in the body. However, it is well established that systemic inflammation in general^3,8^ and PGE2 i.c.v. in particular^77^, promote corticosterone release in rodents. Therefore, elevated corticosterone levels triggered by PGE2 could also function as a stress signal from the body to the brain, suggesting caution when interpreting the results of this work as purely central.

The profound effect of PGE2 i.c.v. on sickness behavior and activation of CAN areas prompted our use of this methodology to explore changes in neuronal activity in neocortical regions previously associated with the representation of body states, such as the IC and the SSp. Our first approach focused on the properties of individual IC and SSp neurons during PGE2-triggered sickness. We found a slight but widespread reduction of firing frequency in these neocortical regions during sickness. A previous report has found that LPS i.p. increases ongoing activity in various cortical areas in anesthetised mice^78^. While differences in experimental conditions, such as recordings in awake versus anesthetised mice, could account for this discrepancy, the absence of additional immune mediators in our reductionist approach may have contributed to the lack of cortical hyperexcitability during sickness.

Next, we focused on the spontaneous population activity of IC and SSp. Using a dimensionality reduction approach with CEBRA and a decoding analysis with GLMs, we observed that both IC and SSp contain information about body temperature and sickness behaviors. These findings are consistent with the observed brain-wide representation of spontaneous behaviors and need states (such as hunger and thirst) across the mouse brain^18,79^. The identification of low-dimensional neuronal activity patterns across multiple brain regions, along with coordinated changes in behavioral profile and strong activation of CAN areas by PGE2, provides further evidence that systemic inflammation triggers a distinct brain state^10^. Additionally, across both methodologies, we observed a consistently better representation of body temperature in IC than in SSp, a difference presumably driven by the warming phase of the fever response. Interestingly, previous work has shown that the posterior IC (pIC) encodes skin warming better than SSp, while neurons responding to skin cooling can be detected in both pIC and SSp^80^. Our findings suggest that this preferential representation of warming in IC over SSp may also extend to internal temperatures.

Additionally, we found better decoding performance for GLMs trained on the IC data compared to SSp across all other sickness features, except for eye squinting, where decoding performance on the SSp was substantially better. Interestingly, strong eye squinting has been associated with visceral pain in rodents^65^, and SSp plays a key role in discriminating sensory inputs associated with pain perception^81^. Therefore, the more substantial representation of eye squinting in SSp than in IC might be related to the nature of this spontaneous behavior as a pain readout and the prominent role of the SSp in pain processing.

In contrast, when we combined all sickness readouts (namely body temperature, eye squinting, displacement velocity, and grooming) into a sickness index, we observed a remarkable separation in GLM decoding performance between IC and SSp, with better decoding performance from the IC data in both hemispheres. Furthermore, while the decoding performance for SI and body temperature from SSp data is comparable, the IC shows a better representation of SI than body temperature alone. These findings suggest that ongoing population activity of IC preferentially encodes a unified representation of body state (such as sickness) rather than individual features associated with such states. This further supports the proposed role of the IC in real-time tracking the internal conditions of the body^82^ and emotions^83^, and the generation of interoceptive predictions to guide behavior^75,82^.

Multiple lines of evidence suggest that the state of the immune system is one of the internal conditions of the body monitored by IC. Studies in mice and humans have consistently identified IC as one of the main hubs in the brain that responds to peripheral inflammation^3,10,11,17,51,74^. Functional studies have also shown that IC can store memory-like information about past inflammatory events in peripheral tissues. Activating these insular inflammation-sensitive neurons after the inflammation is resolved can reestablish inflammation in recovered tissues^84^. Additionally, IC is necessary to form associations between certain environments and specific immune responses^85^. This process is known as immune conditioning, and it is believed to be a type of learning that helps the body anticipate the encounter with pathogens and prepare the immune system accordingly^86,87^.

In agreement with the proposed role of IC in immunoception^16,48^, our Fos map also identifies the IC, particularly the left IC, as activated upon PGE2 i.c.v. Our CEBRA and GLM analyses revealed certain lateralization of sickness representation within the IC. Using CEBRA embeddings, we detected a better representation of body temperature and body temperature combined with sickness behaviors in the left IC, whereas sickness behaviors alone were better decoded from the right IC. Conversely, GLM decoders trained on data from the left IC consistently outperformed those trained on the right IC, a pattern also observed for the SSp. Indeed, the differential contribution of the left versus right IC to sickness and immune state representation remains debatable across the literature, with reports supporting bilateral or unilateral (right or left) insular engagement during systemic inflammation^10,17,51,74,84,85^. Further work will be needed to clarify any potential lateralization in sickness representation and immune system regulation in the IC.

Altogether, our work introduces PGE2 i.c.v. as a powerful tool for studying immune-brain communication in awake mice and highlights the IC as a key region for representing preferentially complex and multivariate information linked to body states, including immune system status.

## Acknowledgments

The authors would like to thank Cornelius Gross (EMBL Rome) for his critical input throughout the project and during manuscript preparation. We also thank the personnel at EMBL’s Laboratory Animal Research facility (LAR), particularly Ernesto Cuevas, Lotte Hoffman, Alessandro Grassi, Christian Grossmann, and Vladyslav Solodovnik, for their excellent support in mouse colony maintenance and animal experimentation. Additionally, we would like to thank Beate Neumann and Stefan Terjung from the EMBL’s Advanced Light Microscopy Facility (ALMF) for their training and support in light microscopy. We also thank Renato Alves from EMBL’s Bio-IT for his assistance with software installation. G.B.K. was funded by an EIPOD4 postdoctoral fellowship under Marie Skłodowska Curie Cofund Actions MSCA-COFUND-FP (664726). This work was additionally supported by the European Molecular Biology Laboratory (EMBL) and the Deutsche Forschungsgemeinschaft (DFG, German Research Foundation) – project 458898724 awarded to RP.

## Author contributions

G.B.K. conceived, designed, and performed the research. J.C.B collaborated with Neuropixel recordings and analyzed the electrophysiology data. D.R. and N.R. conducted the Fos stainings and analysis. M.A.H., J.C.B. G.B.K., and A.C. analyzed behavioral data. J.C.B. and M.A.H. performed statistical analysis and prepared the figures. G.B.K. led the project with collaboration from M.Y.A., M.N.H., M.S., N.R., H.A., and R.P.. G.B.K. and R.P. acquired funding and supervised the project. G.B.K. wrote the paper, and all authors commented on and approved the final version of the manuscript.

## Methods

### 1. Key resources table

| REAGENT AND RESOURCE | SOURCE | IDENTIFIER |
| --- | --- | --- |
| **Antibodies** |  |  |
| Rabbit monoclonal anti-cFos | Synaptic System | cat#226-008 |
| **Mouse strains** |  |  |
| C57BL/6J | EMBL's colony and Charles River Laboratories | Strain #:000664<br>RRID: IMSR_JAX:000664 |
| **Chemicals** |  |  |
| Atipamezol hydrochloride | Vetoquinol GmbH |  |
| Clozapine N-oxide | Enzo Life Sciences | cat#BML-NS105-0025 |
| CM-Dil Cell-Labeling Solution | ThermoFisher | cat#V22888 |
| Dimethyl sulfoxide | Sigma Aldrich | cat#D2650-5X5ML |
| Fentanyl citrate | Piramal Critical Care Deutschland GmbH | PZN#13624067 |
| Flumazenil | CHEPLAPHARM Arzneimittel GmbH | PZN#03488574 |
| Glutaraldehyde solution | Merck Sigma | cat#340855-25ML |
| Isoflurane | Baxter |  |
| Lidocaine hydrochloride (Xylocaine) | Aspen Germany GmbH | PZN#01138002 |
| Lipopolysaccharides from Escherichia coli O111:B4 (LPS) | Sigma Aldrich | cat#L2630-10MG |
| Medetomidine hydrochloride | CP-Pharma | cat#798-491 |
| Meloxicam | Boehringer Ingelheim | cat#796-617 |
|  | Vetmedica GmbH |  |
| Midazolam | CHEPLAPHARM<br>Arzneimittel GmbH | PZN#03085793 |
| Paraformaldehyde solution 4% in PBS | ThermoScientific | cat#30525-89-4 |
| PGE2 | STEMCELL<br>Technologies | cat#72194 and 72192 |
| Saline sterile | Braun | PZN#02737756 |
| Sodium azide 0.05% | Merck Sigma | cat#RTC000068-1L |
| Tergazyme | Alconox | cat#1304-1 |
| Xylocaine (Lidocaine hydrochloride) | Aspen Germany GmbH | PZN#01138002 |
| **Software** |  |  |
| Matlab | Mathworks | 2024a,b |
| Bonsai | https://bonsai-rx.org/ |  |
| SHARP-Track | https://github.com/cortex-lab/allenCCF/tree/master/SHARP-Track |  |
| Kilosort4 | https://github.com/MouseLand/Kilosort | Version 4 |
| FaceMap | https://github.com/MouseLand/facemap |  |
| Boris | https://www.boris.unito.it/ |  |
| CEBRA | https://github.com/AdaptiveMotorControlLab/cebra | 0.4.0 |

### 2. Animal Use Ethics and Maintenance

Experimental procedures on living animals followed the European Union Council Directive 2010/63/EU on the protection of animals used in scientific research. EMBL’s Institutional Animal Care and Use Committee (IACUC) reviewed and approved under license number 22-005_HD_RP all animal experiments in this work. Mice of both genders were obtained from EMBL’s colonies and Charles River Laboratories. Animals were housed in groups of 1-5 in Makrolon type 2L cages on ventilated racks at room temperature and 50% humidity with a 12 hr light cycle and ad libitum access to food and water unless otherwise specified.

### 3. Surgical procedures

#### 3.1 Pre-incisional stereotaxic procedures

All surgical procedures were conducted in aseptic conditions. 8-12 weeks-old mice of both genders were anesthetized with a 5 µl/g intraperitoneal injection of a mixture of Fentanyl (0.07 µg/g), Midazolam (14 µg/g), and Medetomidine (1 µg/g) (aka FMM). Anesthetized mice were transferred to a digital stereotaxic apparatus (cat# 68018; RWD). The head was carefully fixed to the setup using a snout mask, a bite bar, and ear bars. Eyes were protected with hydrating ointment (Bepanthen with 5% dexpanthenol; Bayer), and body temperature was monitored and maintained using a rectal probe and a warm blanket connected to a homeostatic monitoring system (cat#69027; ThermoStar, RWD). Hair from the head was removed with depilatory cream (Veet; Reckitt). Mice received a 5 µl/g subcutaneous injection of Meloxcan (1 µg/g) in sterile saline for postoperative analgesia. Before the incision, the head’s skin was disinfected using alcohol pads (cat#9160612; Braun), and mice received a 50 µl subcutaneous injection of Xylocaine 1% (PZN#01138002; Aspen Germany GmbH) under the skin of the scalp. The skin of the scalp was removed using fine scissors (cat#15006-09; Fine Scientific Tools).

#### 3.2 Brain cannula and head bar implantations

All surgical procedures were conducted in aseptic conditions. Two small craniotomies of ∼⌀1mm were drilled using a microdrill (cat#78001; RWD). One craniotomy was drilled over the lateral ventricle (from bregma AP=0.00 mm, ML=+/-0.80 mm), and the second craniotomy was drilled over the parietal bone. The exposed dura was carefully torn with fine forceps (Dumont#5; Fine Science Tools). A sterilized stainless steel surgical screw (cat# 00-96×1/16, Plastic One) was fixed to the skull over the parietal craniotomy, and a sterilized i.c.v. guide cannula (OD 0.48x ID 0.34x L3 mm; cat#62003; RWD) was carefully lowered onto the brain through the craniotomy over the lateral ventricle. When required, a custom-made head bar was additionally placed over the skull. The cannula’s pedestal, head bar, surgical screw, exposed skull, and skin on the head were secured with glue (Cyano retarder; Hager Werken) and dental cement (Paladur; Heraeus). Once the glue was set, a sterilized dust cap (O.D. 0.30 mm/M3.5, length 2.5mm; cat#62102; RWD) was crewed on the guide cannula to keep the cannula clean and to prevent dirt from entering the brain.

#### 3.3 Logger implantation

All surgical procedures were conducted in aseptic conditions. After stereotaxic procedures, mice were removed from the stereotaxic frame and were carefully placed in a supine position over a warm blanket connected to a homeostatic monitoring system (cat#69027; ThermoStar, RWD). Hair on the belly was removed with depilatory cream (Veet; Reckitt), and the exposed skin was disinfected with alcohol pads (cat#9160612; Braun). A small incision of ∼1 cm was performed on the skin of the abdomen. A sterile implantable temperature logger (DST nano-T; Star::Oddi) was inserted into the abdominal cavity, and muscle and skin layers were sutured separately with sterile absorbable surgical threads (Marlin 17241041; Catgut, Germany).

#### 3.4 Craniotomies for Neuropixel probes

Two small craniotomies of ∼⌀1mm were drilled over the right and left insular cortices (from bregma AP=1.00 mm, ML=+/-3.25 mm) using a microdrill (cat#78001; RWD), and the exposed dura was carefully torn with fine forceps (Dumont#5; Fine Science Tools). After that, the craniotomies were covered with an absorbable hemostatic gelatin sponge (Equispon; Invotec) embedded in sterile cortex buffer (mM: NaCl 125, KCl 5, Glucose 10, Hepes 10, CaCl_2_ 2, MgCl_2_ 2, pH 7.4) and sealed with non-toxic silicon glue (Kwik-Cast; World Precision Instruments).

#### 3.5 Recovery from FMM anesthesia and postsurgical care

FMM anesthesia was antagonized with a 10 µl/g subcutaneous injection of Flumazenil (0.74 µg/g) and Antipamezole (2.8 µg/g). Mice were transferred to their home cage and kept overnight over a heating pad. The following day, mice received a 5 µl/g subcutaneous injection of Meloxcan (1 µg/g) in sterile saline for pain management, and wound healing was closely monitored. Mice carrying brain cannulas were single-housed to prevent cage mates from nibbling on cannula pedestals.

### 4. Training on the treadmill apparatus

This training aimed to habituate mice to the experimenters and to walk on the treadmill on the Neuropixel setup. The experimenter gently handled mice for ∼5 to 15 minutes/day. While handling, mice were offered a sweetened soy drink (cat#4337185832758; Take it veggie) at random intervals as a reward using a Pasteur pipette. After handling, the mice were returned to their home cage. This step was repeated between 3 to 7 days, depending on mouse habituation performance.

Next, mice were briefly anesthetized with isoflurane 5% in O_2_ and head-fixed to the custom-made treadmill apparatus using the previously implanted head bar. Once in the behavioral setup, mice remained on the treadmill between 15 and 60 minutes/day, depending on their habituation progress. While head-fixed, mice were offered a sweetened soy drink (cat#4337185832758; Take it veggie) at random intervals using a Pasteur pipette. After habituation, mice were returned to their home cage. This step was repeated for 3 to 10 days, depending on habituation progress.

### 5. Electrophysiological recordings

24 hours after Neuropixel craniotomy drilling, mice were anesthetized with isofluorane 5% in O_2_ and transferred to the stereotaxic apparatus, where 1-2% isofluorane in O_2_ was vaporized through a snout mask at a flow rate of 0.4-0.6 LPM during surgical procedures. The silicon sealant was carefully removed, and exposed insular craniotomies were covered in sterile cortex buffer (mM: NaCl 125, KCl 5, Glucose 10, Hepes 10, CaCl_2_ 2, MgCl_2_ 2, pH 7.4). While still anesthetized, mice were head-fixed to a custom-made treadmill apparatus and transported to the recording setup. The cap covering the guide cannula was unscrewed, and an i.c.v. injection needle (O.D.0.30mm-with M3.5 G1:0.5MM; cat#62203; RWD) was lowered through the guide cannula. Two Neuropixel 1.0 probes^60^ with their shanks coated in CM-DiI Cell-Labeling Solution (cat#V22888; ThermoFisher), were lowered through the insular craniotomies at ∼10 µm/s using micromanipulators (SMX single-axis; Sensapex piezo-driven micromanipulator system and MPC-385; Sutter motorized micromanipulator system). The electrodes were allowed to settle for ∼15 minutes before starting the recordings. Neuropixel recordings were acquired for 60 minutes, as detailed before^88^. Synchronized with Neuropixel data acquisition, body posture, face conformation, and pupil diameter were monitored and recorded using two IR-sensitive cameras equipped with a CMOS OV2710 sensor (cat#USBFHD05MT-KL36IR; ELP) at 30 fps and 720p resolution under IR illumination (cat#B0BJFGF2YP; Serlium). Synchronization between IR cameras and neuropixels acquisition was performed using Bonsai (https://bonsai-rx.org/) and an Arduino Uno board to capture the cameras’ recording start signal through a digital channel of the neuropixels acquisition system through an NI DAQ board (National Instruments, PXIe-6341).

Ten minutes into the recordings, 2 µl of a sterile solution of PGE2 2 nmol/µl (cat#72194 and 72192; STEMCELL Technologies) in DMSO 1:1 (cat#D2650-5X5ML; Sigma Aldrich) or vehicle (sterile saline and DMSO 1:1) were injected in the lateral ventricle at 1 µl/minute using the previously placed i.c.v. injection needle. After i.c.v. perfusion, electrophysiological recordings were allowed to proceed for another 50 minutes. Upon experiment completion, the Neuropixel probes and i.c.v. injection needle were removed from the brain. Mice were released from the treadmill apparatus, transported to an euthanasia chamber, and sacrificed by CO_2_ inhalation. Euthanized mice were transcardially perfused with BPSx1 followed by PFA 4% in PBSx1 (cat#30525-89-4; ThermoFisher). Brains were collected for electrode track reconstruction.

### 6. Behavioral recordings in freely moving mice

#### 6.1 Recording protocol

Before behavioral recordings, mice were food-restricted for 24 hours. On the day of the experiment, mice were transferred in their home cage to the behavioral room and were allowed to acclimatize for approximately 2 hours before any further procedures. After the injection of PGE2 or LPS, mice were returned to their home cage, where their behavior was monitored using a USB camera (2K Webcam Auto Focus Ultra HD). The top of the home cage was replaced by an acrylic frame to gain better visual access. Camera acquisition was controlled using Bonsai (https://bonsai-rx.org/). Behavior was monitored for up to 2hs (PGE2 injections) or 5hs (LPS injections). After video recordings, mice were transferred back to the holding room.

#### 6.2 LPS injection

Mice were injected intraperitoneally with ∼200 µl of 0.02 mg/kg of LPS in sterile saline^89^ or an equal volume of saline.

#### 6.3 PGE2 injection

Mice were restrained using a disposable restrainer (DecapiCones; Braintree). The pedestal of the guide cannula was pulled out of the restrainer using a small hole cut in the upper part of the restrainer. The cannula’s cap was unscrewed, and a sterile i.c.v. injection needle (O.D.0.30mm-with M3.5 G1:0.5MM; cat#62203; RWD) was lowered through the guide cannula. 2 µl of a sterile solution of PGE2 2 nmol/µl (cat#72194 and 72192; STEMCELL Technologies) in DMSO 1:1 (cat#D2650-5X5ML; Sigma Aldrich) or vehicle (sterile saline and DMSO 1:1)^33^ were injected in the lateral ventricle at 1 µl/minute using a Hamilton syringe (cat#HAM80075; Hamilton) connected with PE tubing (cat#62329; RWD). The injection needle was kept in place for two minutes before retraction to prevent liquid backflow. After infusion, the cannula cap was screwed back, and mice were released from the restrainer and returned to their home cage.

### 7. Histological procedures

#### 7.1 Fos iDisco

Mice of both genders were used in this experiment. I.c.v. cannulas and body temperature loggers were implanted, as mentioned above. Treated mice received 4 nmol PGE2 in DMSO 50% i.c.v. or an equal volume of vehicle (i.c.v. injection procedures detailed above). 1.5h after i.c.v. injection mice were transported to an euthanasia chamber and sacrificed by CO_2_ inhalation. Euthanized mice were transcardially perfused with BPSx1 followed by PFA 4% in PBSx1 (cat#30525-89-4; ThermoFisher). Brains were collected and postfixed in PFA4% at 4°C overnight. Brains were rinsed in PBS1x and stored in PBS1x with azide 0.05% (cat#RTC000068-1L; Merck) at 4°C until further processing.

Fos iDISCO+ immunolabeling was performed as previously described^52^, using a rabbit monoclonal anti-Fos antibody at 1/2000th dilution from the stock (Synaptic Systems, cat#226-008). Brains were imaged on a Blaze light sheet microscope.

#### 7.2 Neuropixel track reconstruction

Brains were post-fixed overnight in PFA 4% in PBSx1 (cat#30525-89-4; ThermoFisher) at 4°C, rinsed in PBSx1, and cryoprotected in sucrose 30% in PBSx1 for a minimum of 24 hs. Brains were embedded and frozen in a sectioning matrix (Tissue-Tek; Sakura). Coronal brain sections were cut at 30 µm in a cryostat (CM 3059S; Leica). Brain sections were collected in glass slides (cat#11976299; Epredia SuperFrost Plus Gold; ThermoFisher) and kept at −20°C until further processing. Slides were rinsed in PBSx1 for 10 min, mounted in DAPI-containing mounting medium (VectaShield Plus; Vector Laboratories), and placed at 4°C in the dark until picture acquisition.

Images were acquired at 4x in an epifluorescence scope (TiE; Olympus) equipped with a Hamamatsu Orca Flash 4 V2 (pixel size:6,5 µm /2048 x 2048 pixel). Neuropixel tracks were reconstructed using the SHARP-Track pipeline^90^.

### 8. Data analysis

#### 8.1 Behavioral analysis of freely-moving mice

Movement data was acquired using pose estimation on video recordings of mouse behavior, recorded at 30 fps. A LightningPose model^40^ was trained to track 26 anatomical landmarks across the animal’s body, including points along the tail, spine, hips, sides, shoulders, ears, eyes, nose, and head center -similar to Ye et al. 2024^91^. After training, videos were processed to generate CSV files containing pose keypoint coordinates and corresponding confidence values. For post-processing, the “movement” package was used^92^. The pose data was filtered by confidence threshold (>0.6), the data interpolated over time (with a maximum gap of 1000 frames), and smoothed with a median filter of 0.1 seconds. We then used the integrated tools of the “movement” package to calculate displacement and velocity.

To correct for perspective distortion and standardize measurements across different recording sessions, we manually annotated the arena boundaries for each recording and applied a perspective transformation using OpenCV. The arena boundaries were mapped to a standardized rectangle coordinate system (320×160 pixels), with this transformation applied to all keypoint positions.

Stasis episodes, defined as periods of minimal movement, were identified through position data processing. First, position data was low-pass filtered with a Butterworth filter with a cutoff frequency of 12 Hz. Position change was calculated as the Euclidean distance between consecutive frames, and a rolling sum of movement was computed using a 5-second window. Frames with rolling movement below a threshold of 25 units were classified as stasis. Continuous stasis periods were identified, and episodes shorter than 5 seconds were excluded from the analysis. Importantly, stasis episodes were adjusted to exclude periods of food interaction to avoid misclassification.

Food interaction periods were detected using a deep learning model trained with DeepEthogram^41^. The model achieved an accuracy of 0.88, with an F1 score of 0.705 and AUROC of 0.938 Figure S1A and B. Using an optimal threshold of 0.633, the model provided a precision of 0.611 and recall of 0.834 for food interaction detection. The mean average precision (mAP) was 0.707. Food interaction prediction confidence scores were thresholded at this optimal value to generate binary classifications. The cumulative time spent interacting with food was calculated for each recording and used for subsequent analysis.

Statistical comparisons between treatment and control groups were performed using multiple approaches. Kolmogorov-Smirnov tests were used to compare non-normal distributions of episode lengths, velocities, and temperatures between treatment groups. Linear mixed effects models assessed treatment effects on cumulative displacement and food interaction over time, incorporating individual mice as random effects to account for repeated measures (modeled as: predictor ∼ injection_consolidated * duration + (1|individuals)). Paired t-tests were used for within-subject comparisons of maximum temperature and cumulative metrics. Analyses were conducted separately for the LPS and PGE2 experiments. Data processing and analysis were implemented using Python and R.

#### 8.2 Behavioral analysis of head-fixed mice

Video data from head-fixed mice obtained during neuropixels recordings were analyzed with point tracking FaceMap^93^ to quantify the % of eye opening as the difference between the y position of a point placed on the top eyelid and another at the bottom eyelid. Treadmill velocity was estimated from a contrasting bar pattern etched on the edge of the disk treadmill and captured in the videos. Treadmill velocity was calculated from simple kymographs of these moving bars obtained in ImageJ. Grooming events were evaluated on a per-second basis manually, assisted by BORIS (https://www.boris.unito.it/). Due to the sparse occurrence of grooming events, grooming frequency was estimated as the per-second number of grooming events averaged over 5 minutes.

Statistical comparisons of the head-fixed behavioral readouts typically consisted of 2-sample Kolmogorov-Smirnov tests between the pooled data from vehicle and PGE2-injected mice, performed using the kstest2 function for MATLAB.

##### 8.2.1 Calculation of a sickness index

To produce an integrated, unidimensional readout summarizing body temperature and head-fixed behavioral dynamics positively correlated to sickness, consisting of -(treadmill speed), -(eye opening), and grooming frequency, we defined a sickness index. To calculate the sickness index, we first normalized the body temperature and behavioral readouts to be bounded between 0 and 1. Then, we defined the sickness index as the sum of these four normalized variables, which were normalized again to range between 0 and 1.

#### 8.3 Neuropixels data processing and analysis

High-pass filtered and common average referenced spike recordings were produced with CatGT (https://billkarsh.github.io/SpikeGLX/#catgt) as described earlier^88^. These preprocessed datasets were spike-sorted and drift-corrected using kilosort4^94^. The output of kilosort was subject to manual curation using Phy2 (https://github.com/cortex-lab/phy), obtaining single-units with somatic AP waveforms, good waveform consistency (amplitude) across spikes, showing few or no refractory period violations (2ms refractory period violation time window) in their correlogram plots and firing a minimum of 400 times in 60 min of recording.

##### 8.3.1 Decoding analysis

For decoding analysis of population activity data we calculated the firing rates of the simultaneously recorded single unit populations as the spike count per 1s time bins. To avoid an influence of the i.c.v. injection manipulation, we excluded from the decoding analysis the 10 minutes pre-injection and the 2 minutes during injection, obtaining 48 minute long firing rate datasets (2880 time points).

To compare the recorded population dynamics in vehicle and PGE2-treated mice, we resorted to the multisession CEBRA-behavior algorithm (https://github.com/AdaptiveMotorControlLab/cebra)^72^ to embed the aforementioned firing rates datasets recorded in each mouse (session) in a common coordinate system through supervised dimensionality reduction guided by accessory variables. The accessory variables we used to supervise the embeddings were: 1) body temperature, 2) % of eye opening, treadmill speed, and grooming frequency simultaneously, or 4) body temperature, % of eye opening, treadmill speed, and grooming frequency simultaneously. After this supervised embedding, we performed cross-validated binary classification of the embedded points by fitting support vector machine (SVM) classifiers to 80% of the embedded firing rate data points and predicting the corresponding treatment (PGE2 or Vehicle) on the 20% left out. We fitted the SVMs using the fitcsvm function from MATLAB with default parameters and evaluated the hit rates as the proportion of correctly predicted labels. Chance-level hit rates were obtained by scrambling the treatment labels and repeating the decoding analysis as described.

Alternatively, we decoded the corresponding body temperature, behavioral readouts, or the sickness index by fitting generalized linear models (GLM) to the firing rate datasets, leaving out every 5th time point for prediction (80/20% cross-validation). We fitted the models to simultaneously acquired data from each mouse during one experiment session, avoiding pooling, using the fitglm function from MATLAB with default parameters. For decoding during body temperature up and down phases, we used a 76/33% cross-validation ratio (one every 3 time points was left out for prediction) to compensate for the roughly half-sized datasets and preserve statistical power for comparing decoding errors. With the fitted models, we predicted the body temperature, behavioral readouts, or sickness index from the time points left out and calculated the prediction error as the difference between the actual values and the predicted values. Since the number of single units simultaneously recorded with one electrode from SSp was typically much larger than in IC, for comparisons between decoding performance at IC and SSp we randomly subsampled the SSp units to decode SSp subsamples of equal size than the collected IC sample. This process was iterated 30 times to better represent the simultaneously recorded SSp population. Chance-level decoding performances were obtained by scrambling the temperature, behavioral, or SI traces and repeating the decoding analysis as described.

Statistical comparisons of the decoding errors or hit rates consisted of either Wilcoxon rank sum or signed rank tests, 2-sample Kolmogorov-Smirnov tests, performed using the ranksum, signrank, and kstest2 functions for MATLAB, followed by Bonferroni’s correction of the obtained p values.

#### 8.4 Fos+ cell density analysis

ClearMap 2^95^ was used for the segmentation and analysis of Fos+ cell distributions from the light sheet scans, as described in (Renier *et al.* 2016)^53^.

## Data availability statement

All raw and processed data supporting this work will be made available upon publication.

## Generative AI and AI-assisted technologies statement

The authors acknowledge the use of Grammarly and ChatGPT to check English grammar and language fluency during the preparation of this manuscript. All authors reviewed and edited the text after employing these tools and are fully responsible for the content of this manuscript.

